# Peptide YY: a novel Paneth cell antimicrobial peptide that maintains fungal commensalism

**DOI:** 10.1101/2020.05.15.096875

**Authors:** Joseph F. Pierre, Diana La Torre, Ashley Sidebottom, Amal Kambal, Xiaorong Zhu, Yun Tao, Candace M. Cham, Ling Wang, Katharine G. Harris, Olga Zaborina, John Alverdy, Herbert Herzog, Jessica Witchley, Suzanne M. Noble, Vanessa Leone, Eugene B. Chang

**Author notes:** **Corresponding Author:** Eugene B. Chang, MD, Martin Boyer Professor of Medicine, Department of Medicine, Knapp Center for Biomedical Discovery, University of Chicago, Chicago, IL.

## Abstract

Perturbed interactions between the intestinal microbes and host correlate with emergence of fungal virulence. Here we report a previously unknown role for peptide YY (PYY), a described endocrine molecule, as an antimicrobial peptide (AMP) expressed by gut immune epithelial Paneth Cells (PC). PC-PYY differs from other AMPs, including lysozyme, because of limited antibacterial activity, packaging in discrete secretory granules, and selective antifungal activity to virulent hyphae, but not yeast forms of *Candida albicans*. The latter action is through binding of cationic PC-PYY to the anionic hyphal surface, resulting in membrane disruption and killing. PC-PYY is compartmentalized to surface mucus, which optimizes activity and prevents conversion to endocrine PYY by dipeptidyl peptidase-IV (DPP-IV). We conclude PC-PYY is a unique AMP with selective antifungal activity that maintains gut fungal commensalism. Compromised PC-PYY action from PC dysfunction and/or mucus depletion in ileal Crohn’s disease may initiate or contribute to disease via fungal pathogenesis.

**Highlights:**

⍰ Paneth Cell PYY (PC-PYY) is an antimicrobial peptide that differs from endocrine-PYY
⍰ PC-PYY is a selective anti-fungal peptide, targeting the virulent form of C. *albicans*
⍰ PC-PYY is separately packaged, retained by mucus, and released by *C. albicans* hyphae
⍰ PC-PYY is proposed as essential for maintenance of fungal commensalism in the gut

**Graphical Abstract:** Model for Paneth cell (PC) PYY action and regulation of fungal commensalisms and potential role in the pathogenesis of ileal Crohn’s Disease (iCD)
**(A)** In a healthy ileum, commensal yeast reside and do not stimulate PYY_1-36_ release from PCs. **(B)** Increased virulent hyphae (purple hyphae) results in PYY_1-36_ release from crypt PCs into the mucus. Hyphae are targeted by PYY_1-36_ and killed (red hyphae) to manage the increased fungi community in gut. **(C)** In a diseased ileum such as iCD, hyphal load induces immune activation and increased inflammation through PC dysfunction (gray PCs) and decreased PYY_1-36_ release or mucus depletion and PC dysfunction.

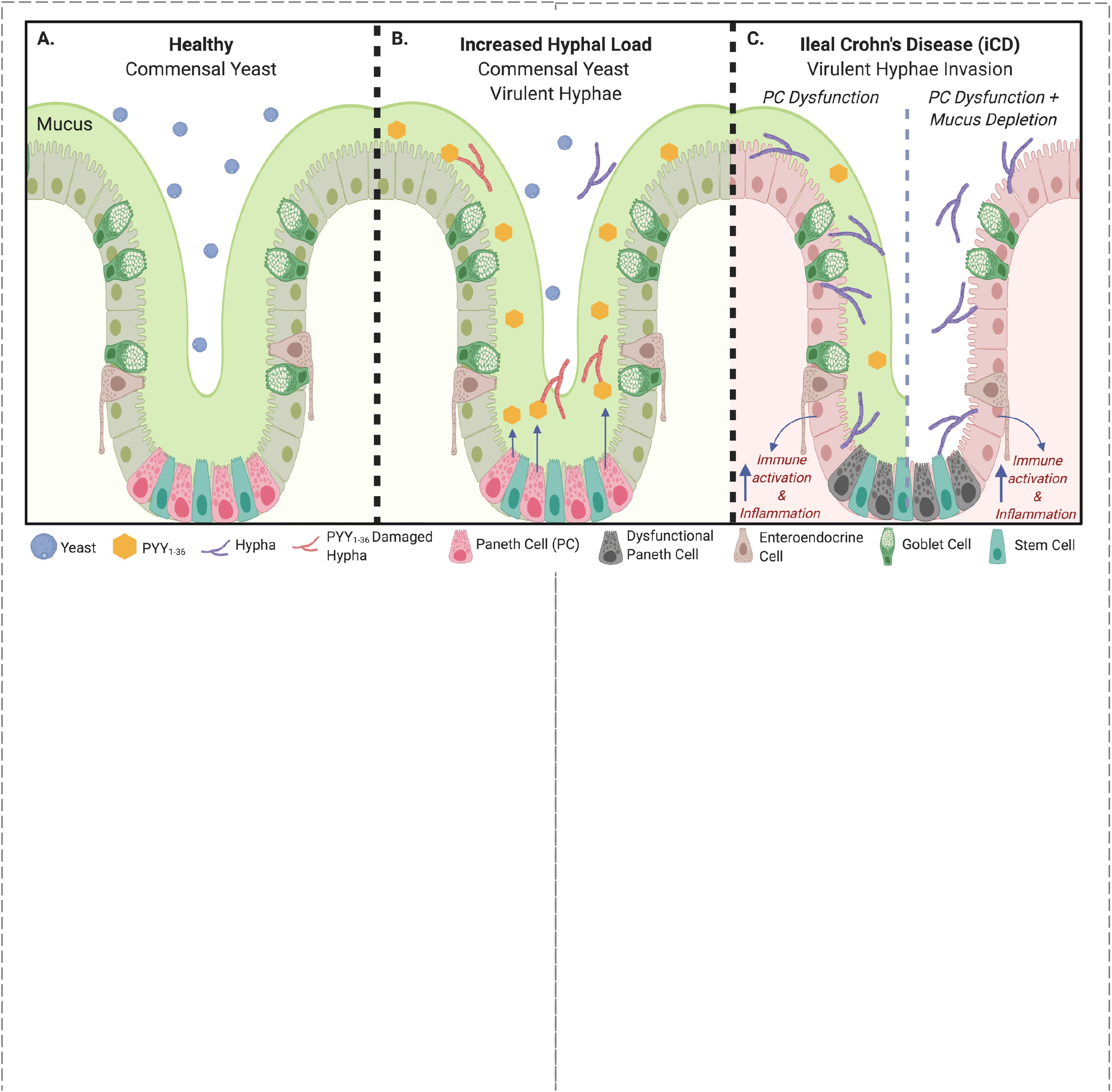

## INTRODUCTION

The coevolution of metazoans and microbes has been paramount to the development of mutualistic beneficial relationships, particularly in the digestive tract where trillions of microbes form region-specific stable and resilient microbial communities essential for processes such as immune and metabolic development and intestinal homeostasis (Costello et al., 2009; Lee et al., 2013; Rakoff-Nahoum et al., 2004). While much of this knowledge has been gained through studies of the Bacteria domain, other microbes such as Fungi and Archaea cohabitate this space. Collectively, these microbes are carefully selected and “tolerated” by the host. This being said, far less is known about how these populations are surveyed and maintained by the host. For fungi, several studies have identified innate immune pathways such as C-type lectins, including Dectin-1 and the mannose receptor, that deter systemic infection through activation of the immune system (Iliev et al., 2012), but their role and actions in deterring fungal infection are not specific and can ultimately cause collateral damage to host cells and other members of the gut microbiome.

Although peptide YY (PYY) is best known as a satiety hormone secreted by enteroendocrine cells (EECs), we now report that PYY is expressed by innate immune Paneth cells of the gut where it appears to be a unique antimicrobial peptide (AMP) with selective activity against the virulent form of polymorphic *Candida albicans. C. albicans* is a commensal yeast of the gastrointestinal tract, but it is also an opportunistic pathogen commonly seen to cause disease in immunocompromised patients (Hoarau et al., 2016; Sokol et al., 2017b). PYY is a highly conserved 36-amino acid peptide produced by diverse vertebrate species that range from cartilaginous fish to humans (Conlon, 2002). Since its discovery in porcine intestine and brain in 1982, PYY has been shown to regulate energy homeostasis and food intake, sympathetic vascular tone, digestion, circadian rhythms, and other endocrine and autonomic functions (Tatemoto, 1982; Batterham et al., 2006; Glavas et al., 2008; Hill et al., 2011; Holzer, 2016). These classically understood endocrine functions of PYY are mediated by the binding of the dipeptidyl peptidase IV (DPP-IV)-modified form of PYY, PYY_3-36_, to the neuropeptide Y receptor (Y2), to which the unmodified form, PYY_1-36_, poorly binds (Batterham et al., 2002; Grandt et al., 1994; Keire et al., 2000). Until our report, intestinal PYY was thought to have solely an endocrine function, released postprandially into the blood circulation by EECs to activate many regions of the brain important to the control of behavior and satiety.

## RESULTS

### PYY is expressed by intestinal innate immune Paneth cells primarily found in the murine and human distal ileal mucosa

While visualizing L-cells in sectioned murine distal ileum by staining PYY with fluorescence antibodies or fluorescence in situ hybridization (FISH), we made the serendipitous discovery that PYY is also present in cells expressing lysozyme, a classical marker of Paneth cells (Figures 1A and 1B). This unanticipated and previously unreported finding was perplexing, as Paneth cells are innate immune epithelial cells of the gut mucosa known to express and secrete AMPs that defend against pathogens and shape the regional gut microbiome. We also confirmed that PYY immunolocalized to Paneth cells in ileal mucosal biopsies from healthy human adults (Figure S1A). We determined that the immunoreactivity was specific to PYY in Paneth cells by using primary antibody pre-adsorbed with recombinant PYY, which resulted in nearly complete quenching of signal from both L- and Paneth cells (Figure S1B). Additional confirmation was made through high resolution images obtained by stimulated emission depletion (STED) microscopy, which revealed that PYY and lysozyme appear to be packaged into discrete granules (Figures 1C and S2), raising the possibility that they might have different roles, sorting pathways, and regulation (Clevers and Bevins, 2013). To further verify that PYY is produced in Paneth cells, laser capture micro-dissection was performed on sections of the murine ileum to isolate distinct populations of basal crypt cells which would include Paneth cells, and villus tip cells where mature L-cells are found (Figure 1D). Confirmation of distinct villus and base crypt populations was conducted by detection of transcripts of Paneth cell (cryptidin 1), and villus cell (sucrase-isomaltase) markers (Figure 1E), respectively. Quantitative RT-PCR was then performed on total RNA extracted from these populations, revealing robust PYY transcript levels in the basal crypt population in line with other Paneth cell-specific AMP transcripts. Finally, we utilized publicly available databases of single cell RNAseq from murine small intestinal epithelial cells. We identified PYY expression in mature enteroendocrine populations through the marker genes Secretogranin-1, CCK, and GLP1, but also in Paneth cell populations defined by the marker genes Cryptdin 17, ATG16L1, DEFAA5, and Lysozyme. (Figures S3A and S3B) (Haber et al., 2017).

**Figure 1.**
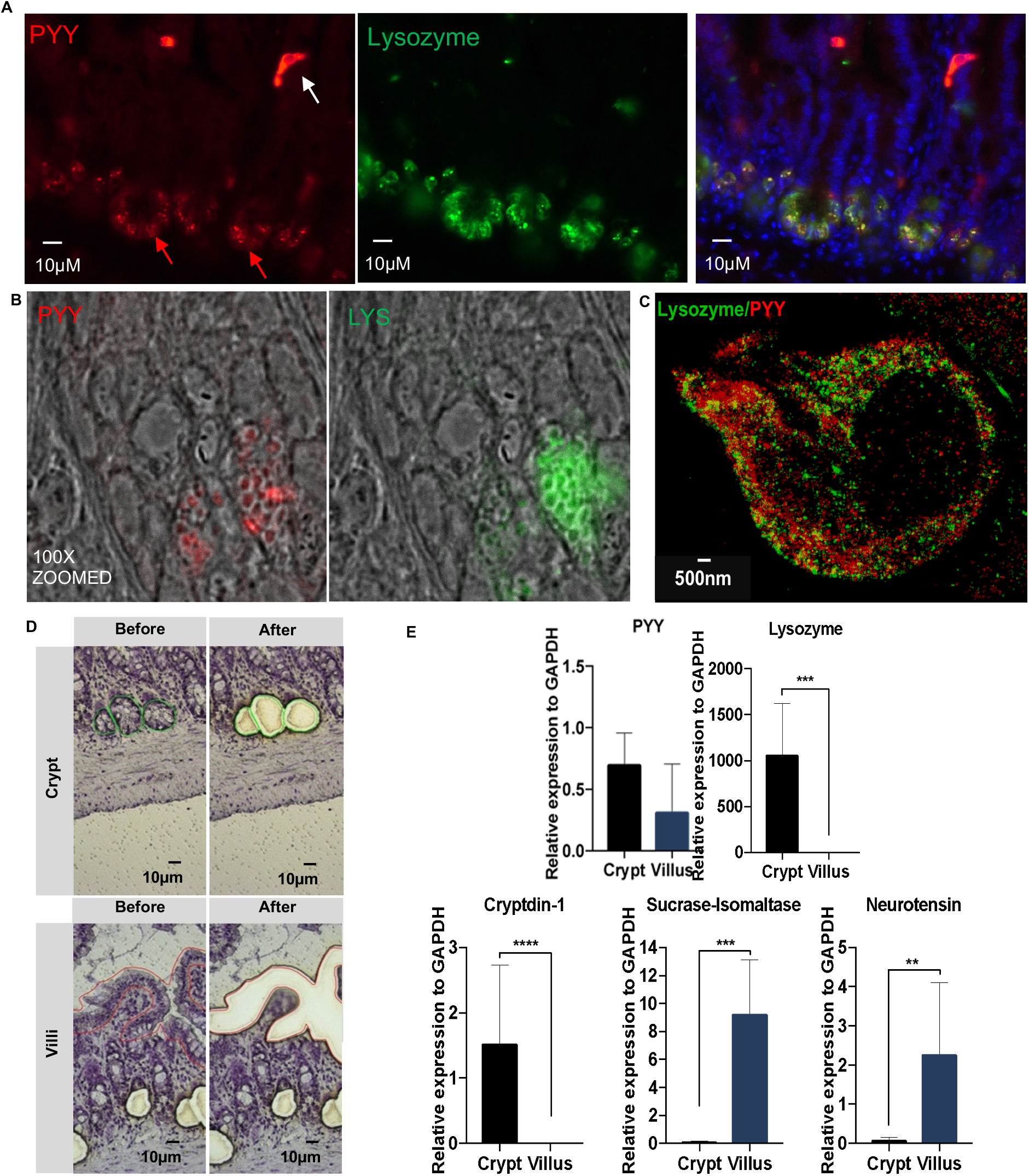
Localization of Peptide YY (PYY) in murine ileal Paneth cells. **(A)** PYY (red) was detected in Paneth cells (red arrow) and L-cells (white arrow) via immunofluorescence. Paneth cell PYY staining co-localized with the Paneth cell enzyme, lysozyme (green; middle panel). **(B)** PYY localization in Paneth cells was was confirmed using PYY mRNA Fluorescent *In Situ* Hybridization (FISH) and counterstaining with lysozyme antibody (green). **(C)** High-resolution stimulated emission depletion (STED) microscopy that indicates immunofluorescence (IF) PYY and lysozyme are separately packaged within distinct Paneth cell secretory granules. **(D)** Laser capture microscopy (LCM) was used to excise mRNA from Paneth cells and villus epithelial cells. **(E)** mRNA expression of PYY was detected in crypt cells and villus epithelium from LCM extractions. Lysozyme and Cryptdin-1 are Paneth cell antimicrobial products. Sucrase-lsomaltase is a brush boarder product found on the villus surface. Neurotensin is an enteroendocrine cell product. The expression of villus and Paneth cell marker genes were consistent with Paneth cell expression at the crypt base in addition to observed expression of PYY in enteroendocrine cells throughout the villus. Significance was measured using t-test (n=6, repeated twice, *p<0.05; **p<0.01; ***p<0.001; ****p<0.0001).

### PYY is a Paneth cell antimicrobial peptide that has limited anti-bacterial activity, but selectively targets virulent, but not commensal *Candida albicans*

The expression of PYY in intestinal Paneth cells, which secretes AMPs into the gut lumen, suggested an alternative antimicrobial function. As many AMPs have previously described broad activity against bacterial species, we assessed the activity of synthesized PYY peptide (amino acids 1-36) for anti-bacterial activity against a panel of representative grampositive and -negative strains. PYY did have significant, but limited effects on *in vitro* viability and proliferation of some Gram positive microbes such as *Lactobacillus rhamnosus, Enterococcus faecalis, Listeria monocytogenes*, and Gram negative *Escherichia coli, Bacteroides fragilis*, but in other cases there was no antimicrobial activity observed, including *Peptostreptococcus anaerobius, Staphylococcus aureus, Salmonella enterica*, and *Pseudomonas aeruginosa* (Figures 2A, 2B and S4). Overall, the anti-bacterial activity of PYY was underwhelming compared to that of other AMPs (e.g., magainin-2, an amphibian analog of PYY, see below) (Figures 2A and 2B). This was surprising because the predicted structure of PYY bears striking resemblance to another alpha-helical, amphipathic AMP, the amphibian analog magainin-2 (Figures 2C and 2D). Magainin-2 is produced in the skin of the African clawed frog (*Xenopus laevis*) where it has a wide spectrum of antimicrobial activity against many species of bacteria and fungi. Because fungi were not included in our initial screen, we then tested the activity of PYY against *C. albicans* (Figure 3). *Candida* species can exist in a commensal yeast state or a virulent multicellular, filamentous hyphal form that includes altered gene expression and functional properties, the latter including mucosal invasion, immune activation, and inflammation (Moyes et al., 2010; Noble et al., 2017)

**Figure 2.**
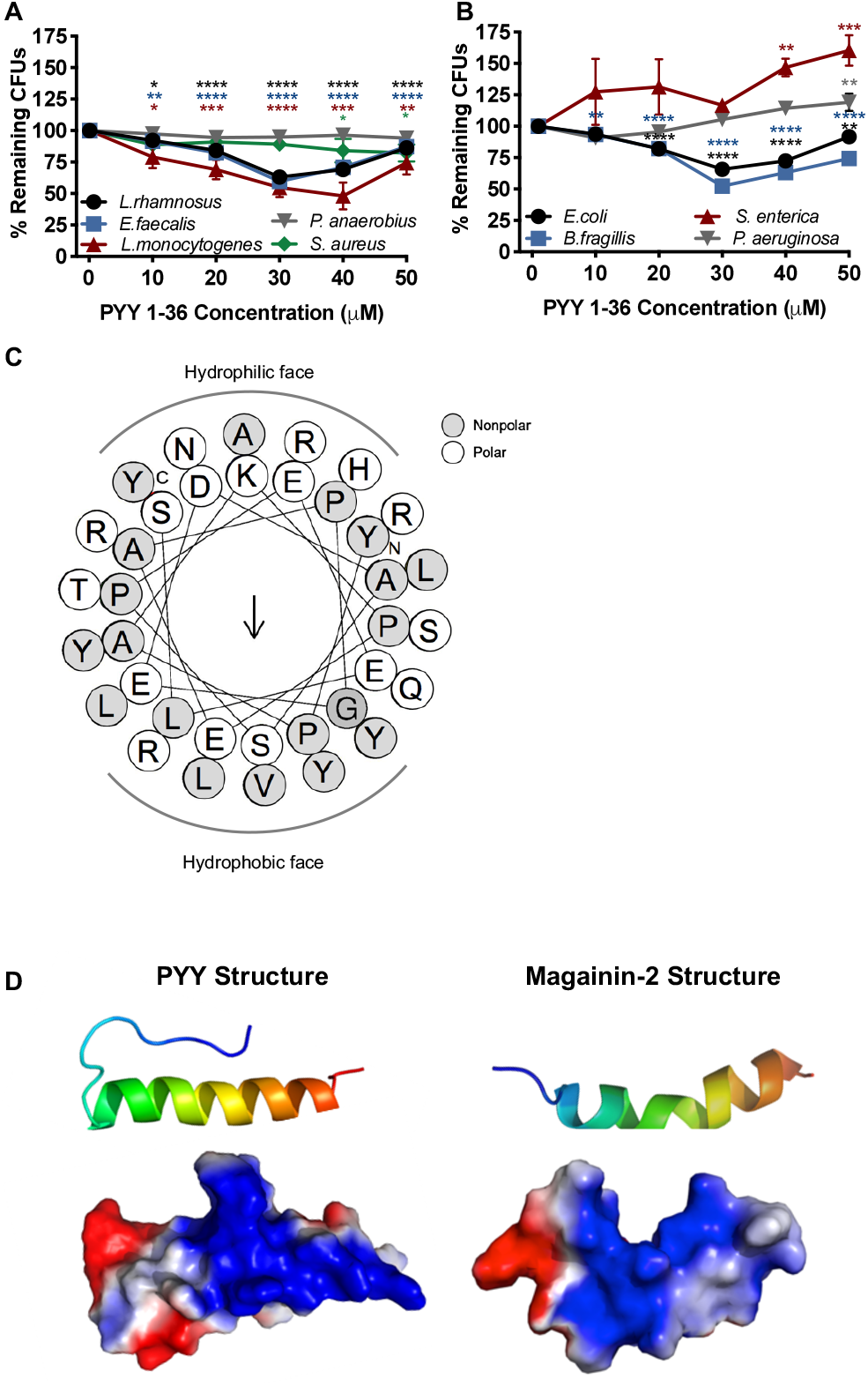
PYY antimicrobial selectivity against bacterial strains and alpha-helix characteristics. Exposure to varying concentrations of PYY for 2 hours results in dose response to killing **(A)** Gram positive and **(B)** Gram negative strains. All assays performed in duplicate three times, CFU values were normalized to the plate containing 0 μM PYY and significance was analyzed by ANOVA (*p<0.05; **p<0.01; ***p<0.001; ****p<0.0001). **(C)** Amphipathic PYY_1-36_ helical wheel projection displaying surface localization of residues and calculated hydrophobic dipole moment (μH 0.208, arrow). **(D)** Ribbon diagram (top) and space-filling model indicating electrostatic surface charge (bottom; red negative, blue positive) for amphipathic alpha-helix structures of PYY_1-36_ and magainin-2.

**Figure 3.**
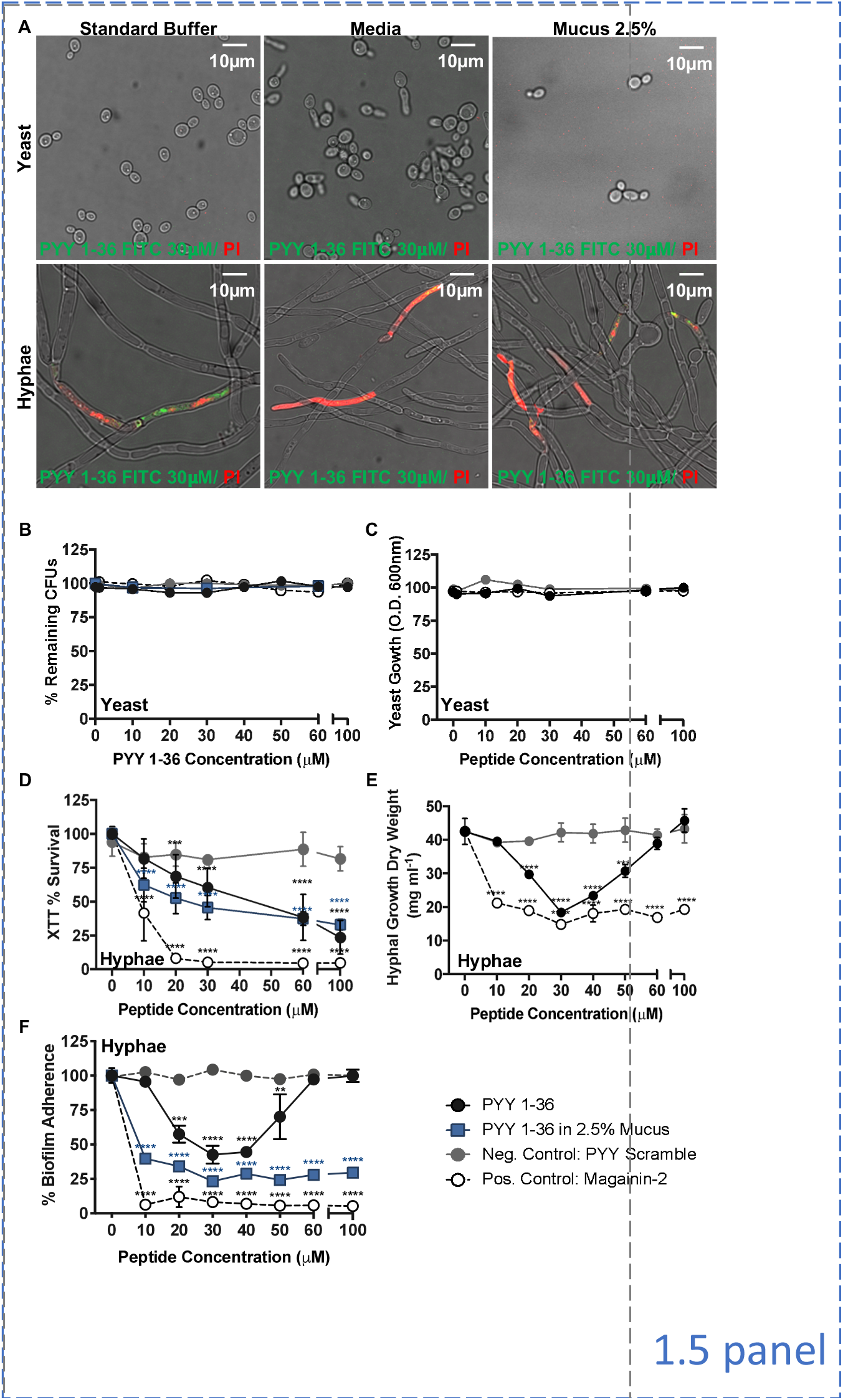
Antifungal activity of PYY_1-36_ against pathogenic *C. albicans* hyphae. **(A)** *C. albicans* were exposed to peptides in the presence of propidium iodide, which enters compromised cells and fluorescently binds DNA. PYY_1-36_ FITC (green) was observed to penetrate only the hyphae form, not the yeast form, of *C. albicans* under several media conditions: standard buffer for AMPs, preferred growth media for each form (YPD-yeast or RPMI-hyphae), and standard buffer with 2.5% w/v porcine mucus. **(B)** Survival as measured by CFUs shows *C. albicans* yeast was not impacted when exposed to PYY at varying doses regardless of the media solution, standard buffer or standard buffer with 2.5% w/v mucus. **(C)** Yeast growth in YPD media evaluated by OD also did not show any decrease with exposure to PYY. **(D)** Conversely, pathogenic *C. albicans* hyphae survival significantly decreased following PYY treatment in standard buffer starting at 20 μm and standard buffer with 2.5% w/v mucus at 10 μm, highlighting the stabilizing effect of mucus on PYY and selectivity of PYY against hyphae. **(E)** Similarly dry weight measurements show PYY inhibits hyphal growth significantly at 20 μm to 50 μm in RPMI media. **(F)** The adherence of *C. albicans* biofilm created by hyphae is also disrupted in RMPI starting at 20 μm to 50 μm and RPMI with 2.5% w/v mucus at 10 μm. RPMI is necessary for hyphal growth and biofilm formation as neither hyphae nor yeast grow in standard buffer, but the bell curve with PYY in RPMI shows the importance of standard buffer media preventing PYY aggregation with increased concentration. All experiments were done in triplicate with 3 replicates, significance was measured using ANOVA and Dunnett’s Multiple Comparisons Test (*p<0.05; **p<0.01; ***p<0.001; ****p<0.0001).

We therefore investigated the effect of PC-PYY (herein also specified as PYY_1-36_) on *C. albicans* growth and viability. As will be shown later, PC-PYY is strictly the unmodified, full length PYY_1-36_, whereas the endocrine form is primarily the PYY_3-36_ form. We used liquid culture conditions which promote the growth of the yeast or hyphal forms of *C. albicans*. Yeast extract Peptone Dextrose (YPD) broth was used to promote yeast growth and RPMI 1640 was used for induction of hyphal growth (Mukaremera et al., 2017; Weerasekera et al., 2016). Additionally, 2.5% w/v porcine mucin (mucus) was included in growth media to investigate environment driven bioactivity within the gut. To determine whether PYY_1-36_ permeabilizes virulent *C. albicans* hyphae, we exposed PYY_1-36_-treated hyphae to propidium iodide (PI), a dye that is excluded from cells with intact plasma membranes and fluorescently labels DNA upon cell entry. PYY_1-36_-treated hyphae are permeabilized at a dose of 30 μM, whereas there is no effect on the yeast form, indicating hyphae-specific bioactivity (Figure 3A). PYY_1-36_ was tested in other species of *Candida*, including *C. tropicalis* and *C. dubliniensis*, where both were susceptible, but the latter was more difficult to assess as the broth conditions were not optimal for yeast to hyphae conversion (data not shown). PYY_1-36_ - fluorescein isothiocyanate (FITC) tagged peptide localizes to the same region of the hyphae that shows PI penetration into the cytoplasm, consistent with direct permeabilization as a mechanism of action.

We further tested the effect of PYY_1-36_ on the viability of *C. albicans* by subjecting both the hyphae and yeast forms to increasing concentrations of PYY_1-36_ in growth media with 2.5% w/v mucus (the natural environment for PC-PYY as will be shown below). PYY_1-36_ showed no effect on yeast viability, as measured with CFUs, but did show significant hyphae death measured with XTT viability in both standard buffer and 2.5% w/v mucus (Figures 3B, 3D and S5). On the other hand, the enteroendocrine form PYY_3-36_ had much less impact on hyphae (Figure S5). The effect of PYY_1-36_ on yeast and hyphae growth was assessed by exposing the two forms in their respective growth media to increasing concentrations of PYY_1-36_, optical density measurements were used for yeast while dry weight measurements were used for hyphae to account for the different properties of the two forms. Again, yeast showed no decrease in growth regardless of PYY_1-36_ concentration, whereas hyphae showed significant growth reduction from 20-50 μM (Figures 3C and 3E). It is notable that the dose-response of PYY_1-36_ action in aqueous (RPMI) buffer is biphasic in functional assays, where the peak effect is ~30 μM with progressive loss of activity by 100 μM. Similar biphasic responses have been reported for other AMPs (Kagan et al., 1994; Phadke et al., 2002), which we attribute to concentration-dependent self-aggregation or multimerization of amphipathic alpha-helical AMPs in aqueous buffer, resulting in decreased bioactivity and bioavailability. Supporting this notion, a typical sigmoidal doseresponse curve is seen when the assay is performed in 2.5% w/v mucus (Figures 3E and 3F). We posit that mucus has properties that prevent self-aggregation of PYY_1-36_ and is the natural environment for PC-PYY that confers optimal activity, retention, and stability (see below), which was supported by luminal vs mucosal mass spectrometry quantification of PYY described below.

As we had observed inhibition of growth and decreased viability of the hyphal, but not yeast, form of *C. albicans* by PYY_1-36_, we next examined whether PYY had an effect on biofilm formation. Hyphal morphogenesis is essential not only for *C. albicans* virulence, but is also required for the establishment of biofilms, which are structured communities of yeasts and hyphae that form on surfaces such as denture material or intravenous catheters. Biofilms promote and exacerbate disease at mucosal surfaces by serving as a source for fungal dissemination and intrinsic resistance to antimicrobial agents. Using *in vitro* assays for biofilm formation that quantitate biofilm by crystal violet staining, we show that PYY_1-36_ in RPMI with or without 2.5% w/v mucus and magainin-2, but not a scrambled peptide control disrupts this clinically significant virulence attribute (Figure 3E and 3F). Again, a classic sigmoidal dose-response curve was observed only when mucus was present.

We next examined the underlying basis for this selectivity for the hyphae form of *C. albicans*. PYY_1-36_ is a cationic, amphipathic alpha-helical peptide with a beta turn and localized negative and positive electrostatic cloud patterns at its surface (Figures 2C and 2D). Other AMPs sharing these structural features are believed to preferentially bind to the anionic surface charge of microbes that then positions the peptide for disruption or pore formation of the lipid membrane that kills or compromises fitness of the microbe (Clevers and Bevins, 2013; van der Weerden et al., 2013). On the other hand, mammalian host cells that have neutral surface charge properties are unaffected by these types of AMPs (Mahlapuu et al., 2016). Using a cationic ferritin probe that identifies anionic surface charge of cell membranes through transmission electron microscopy (TEM), we observed a dramatic difference in surface charge of yeast and hyphae membranes (Figure 4A). Hyphae membranes were found to be anionic, as they were decorated with the cationic probe on their outer surface, whereas the yeast membranes lacked any labeling. Using the same preparation, scanning electron microscopy (SEM) of these two states of *C. albicans* was performed and showed that exposure of *C. albicans* hyphae membranes to PYY_1-36_ was associated with surface blebbing and irregularities, as opposed to the yeast membranes that appeared unaffected (Figure 4B).

**Figure 4.**
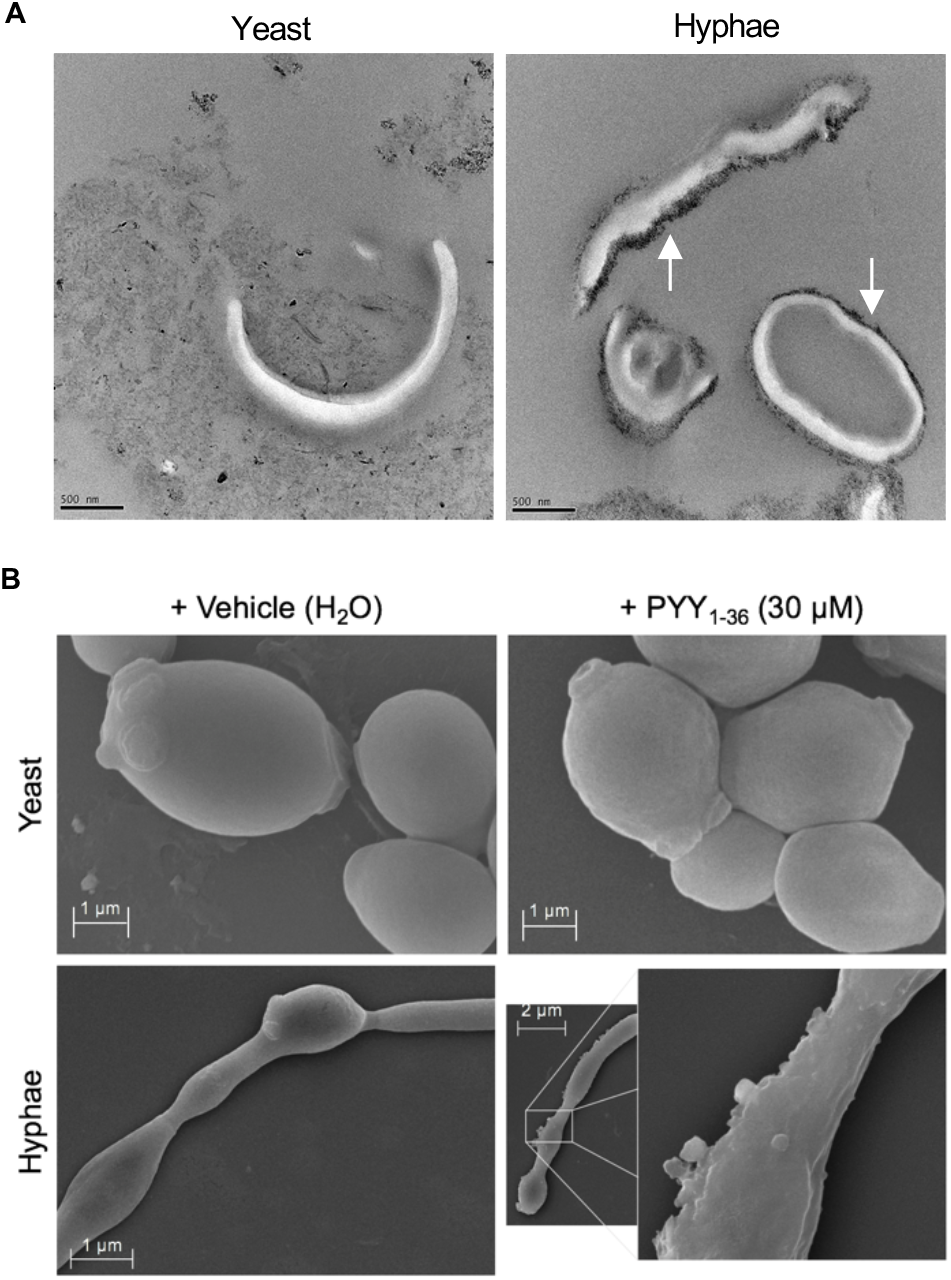
*C. albicans* surface charge and morphological changes associated with exposure to PYY_1-36_. **(A)** *C. albicans* yeast and hyphae membranes were frozen, fractured, and stained with a cationized ferritin probe and examined by a transmission electron microscopy (TEM) for visualization of cationic surface (black probe accumulation). **(B)** *C. albicans* yeast and hyphae forms were treated with vehicle (H_2_O) or PYY_1-36_ (30 μm) for 2 hours and imaged by scanning electron microscopy (SEM). Membrane blebbing and lipid membrane changes were observed in hyphae treated with PYY_1-36_ compared with yeast.

In summary, using five different assay approaches (biofilm formation, propidium iodide uptake, CFU counts, metabolic respiration (XTT), TEM and SEM), we confirm that PYY_1-36_ has selective antimicrobial activity against *C. albicans*, preferentially inhibiting growth and/or killing the hyphal form with little or no activity against the yeast form.

### Paneth Cell PYY (PYY_1-36_): A unique AMP different from Endocrine-PYY (PYY_3-36_)

A hallmark of Paneth cells, which are localized deep in intestinal crypts, is their vectoral secretion of secretory contents such as AMPs into the gut lumen in response to microbial products (Clevers and Bevins, 2013). This is in contrast to PYY of enteroendocrine L-cells that are found in the upper mucosal layers where they secrete PYY and other hormones into mucosal capillaries following food consumption. Although the same PYY_1-36_ form produced by Paneth cells is initial secreted by L-cells, it is rapidly converted by the ubiquitous dipeptidyl peptidase-IV (DPP-IV) to PYY_3-36_, the major circulating endocrine form that binds to the Y2 receptor to mediate PYY’s postprandial endocrine actions (Keire et al., 2010). To examine whether PC-PYY is similar to other PC AMPs in route of action, we used an *ex vivo* preparation where the murine distal ileum is excised, flushed and ligated at both ends, filled with growth media, and maintained at 37 ^o^C (Figure S6). PYY_1-36_ was detected by LC-MS (Figure S7) from the luminal compartment (L), overlying surface mucus (M), and mucosal tissue (T) under basal (no treatment) and treatment conditions, after the mucosa was exposed for 2 hours to *C. albicans* yeast, hyphae, or their respective conditioned media (Figure 5A). Minimal PYY_1-36_ was detected in the luminal (L), overlying mucus (M), and mucosal tissue (T) of *ex vivo* intestinal loops under basal conditions. However, in the presence of *C. albicans* hyphae, but not with yeast or conditioned media from either forms, PYY_1-36_ levels in the mucus compartment increased significantly, whereas no PYY_1-36_ was detectable in luminal compartment under any of the stimulation conditions. Moreover, PYY_3-36_ was not detected in any of the samples, a finding similar to a previous report in rat ileum where PYY_1-36_ was the dominant luminal form (Greeley et al., 1987; Keire et al., 2010). These findings suggested that the PC-PYY is released in response only to the virulent (hyphae) form of *C. albicans* into the overlying mucus where it is retained. As little PYY_3-36_ was detected in tissue, we speculate that it either rapidly disseminated into the blood circulation or was lost during the tissue preparation procedure. We then explored the possibility that PYY_1-36_, because of its cationic amphipathic properties, has proclivity for the hydrogel properties of mucus. An *in vitro* phase-transition partitioning experiment was performed where mucus and aqueous compartments were separated by a 10 kDa MWCO filter (Figure 5B). Regardless of whether the initial placement of PYY_1-36_ was in the mucus or aqueous compartment, it preferentially partitioned into the mucus compartment, providing a plausible explanation as to why PYY_1-36_ is retained in mucus. What remained unexplained, however, was why PYY_1-36_, and not PYY_3-36_ is readily detected in overlying mucus where DPP-IV is present (Darmoul et al., 1994; Michel et al., 2008), a finding we confirmed by immunofluorescence (IF) (Figure 5C). We therefore compared DPP-IV *in vitro* activity to PYY_1-36_ suspended in either aqueous buffer or porcine gastric mucus, assessing the potential resulting PYY products with LC-MS (Figure 5D). In aqueous buffer, DPP-IV converted PYY_1-36_ to PYY_3-36_, and no other proteolytic products were found. However, when performed in mucus, PYY_1-36_ did not undergo DPP-IV proteolysis, suggesting the polypeptide arm of PYY may facilitate mucin binding and therefore be protected from proteolytic cleavage. Alternatively, mucus could be decreasing DPP-IV enzymatic efficiency. Collectively, these data show that secretion of PC-PYY (PYY_1-36_) is preferentially triggered by *C. albicans* hyphae into overlying mucus where it is retained and protected from enzymatic conversion to PYY_3-36_ by DPP-IV, allowing it to function as a unique AMP. These properties and regional compartmentalization could well fortify innate mucus defenses against *C. albicans* hyphae, while not affecting commensal yeast forms or other members of the gut microbiota outside of the mucus barrier and in the lumen.

**Figure 5.**
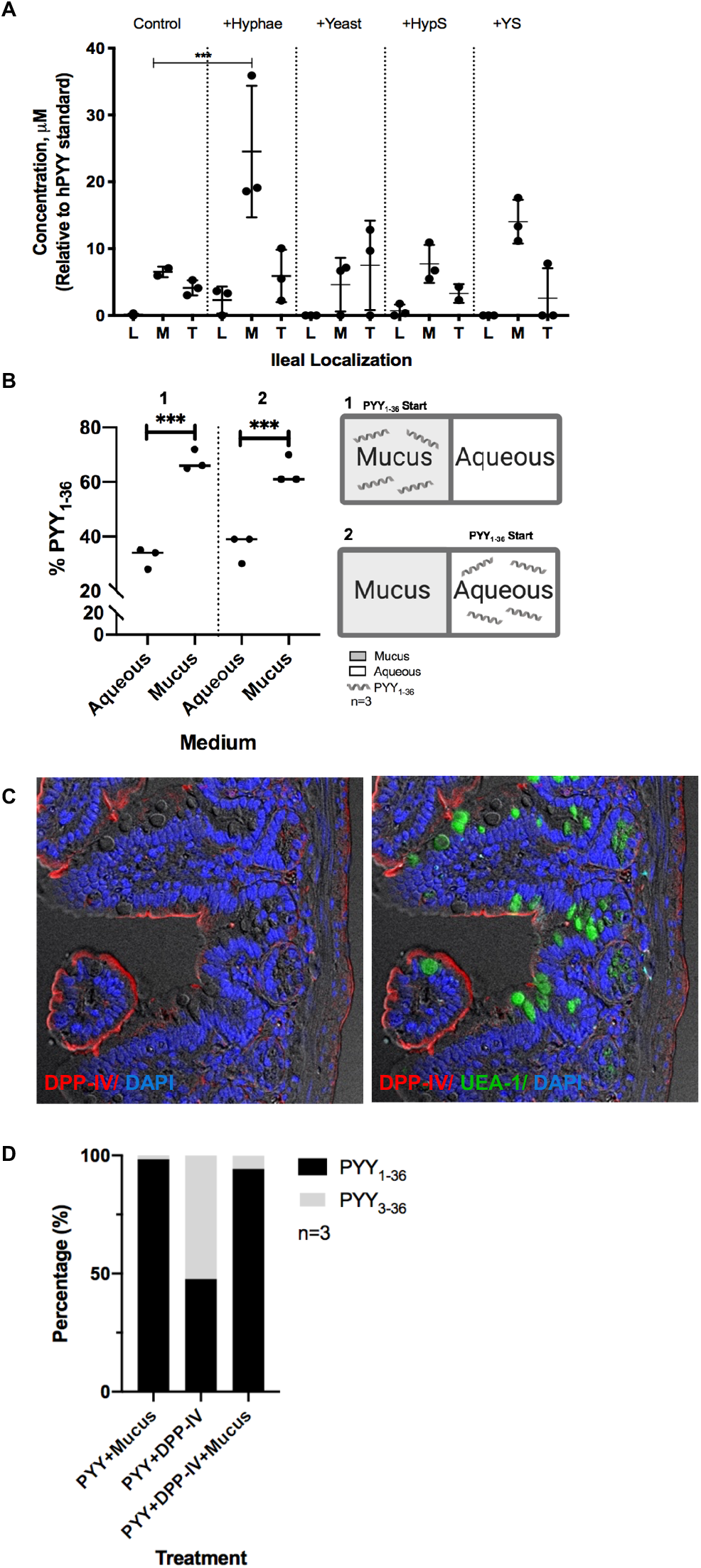
Exposure to *C. albicans* hyphae but not yeasts enhances PYY localization into ileal mucus. **(A)** Quantification of PYY within excised ileal loops following stimulation with vehicle buffer control, *C. albicans* hyphae, *C. albicans* yeasts, hyphae culture supernatant, or yeasts culture supernatant. PYY_1-36_ was quantified in the lumen (L), mucus (M), and tissue (T), where a significant increase in PYY 1-36 was detected when loops were stimulated with hyphae (+Hyphae) compared to all other conditions (+YS: yeast supernatant, +HypS: hyphae supernatant), (n=3, ***One-way ANOVA p<0.001). Ions were detected by liquid chromatography-electrospray ionization-mass spectrometry (LC-ESI-MS). **(B)** PYY partitions to the mucus layer in an *in vitro* partitioning assay. PYY_1-36_ has a higher affinity for mucus compared to aqueous when partitioned between a 10 kDa MWCO filter. Peptide was added to either the mucus (1) or the aqueous (2) and then the amount of peptide on each side was quantified by LC-ESI-MS. **(C)** Immunofluorescence (IF) of paraffin embedded ileal mouse tissue shows that dipeptidyl peptidase IV (DDP-IV) is present around the brush border. Counterstaining was performed with Lectin UEA-I-FITC (*Ulex europaeus*) for ileal mucus and DAPI for nuclei. **(D)** PYY was exposed to DPP-IV under various conditions to assess peptide degradation. Efficiency of DPP-IV cleavage of PYY_1-36_ to PYY_3-36_ was decreased when treated with mucus, as compared to DPP-IV and PYY_1-36_, no mucus. Enzymatic cleavage products were confirmed by LC-ESI-MS (n=3).

### *In vivo* and *in vitro* actions of PYY_1-36_ on *Candida albicans*

We next sought to further determine the correlation of the above findings to human and murine physiology and pathophysiology. To simulate an intestinal environment, we used human colonic epithelial Caco-2 cells that form monolayers and are able to produce mucus. GFP-tagged *C. albicans* grown in a hyphal form were plated over these cells and exposed to PYY_1-36_. The number of attached hyphae were significantly reduced after 4 hours in the presence of PYY_1-_ 36 (Figures 6A and 6B). To determine the impact of PYY on fungal populations *in vivo*, we utilized a mouse model of chronic intestinal colonization in which *C. albicans* persists in the gut lumen and the host remains healthy. The mouse model uses specific pathogen free wild-type (WT) or PYY^-/-^ gene-deficient mice on the C57Bl/6 background. Oral gavage of colonized animals with PYY_1-36_ decreased fungal titers in stool (Figure 6C). Conversely, the rate of fungal colonization in PYY deficient (PYY-KO) mice was more rapid, achieving a level 2-3-fold higher than in wild type control animals (Figure 6D). Of note, total fecal output remained similar in untreated, PYY-treated, and PYY-KO mice, arguing against changes in the rate of fungal excretion as a determinant of fungal load.

**Figure 6.**
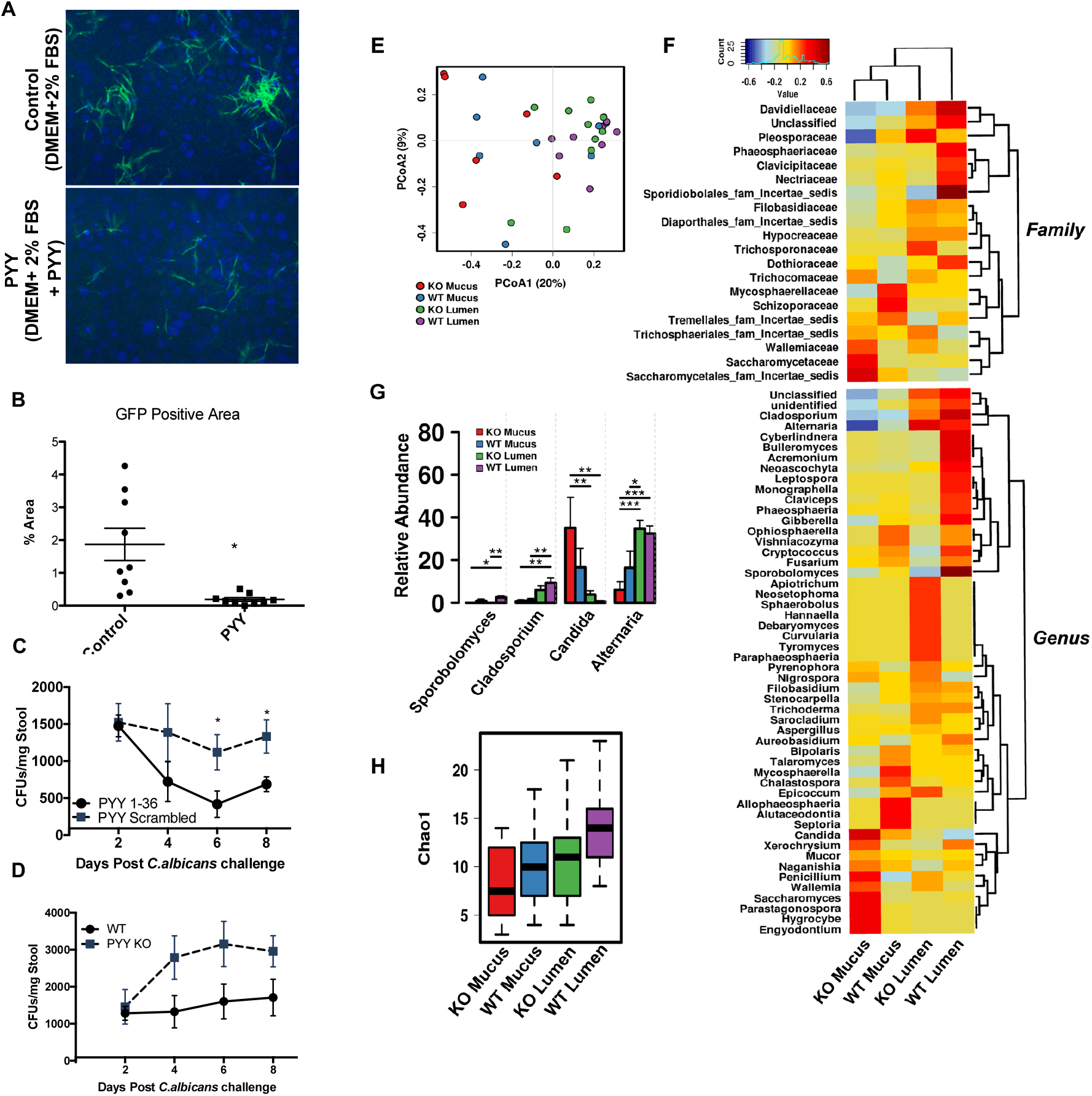
PYY reduces gastrointestinal colonization of *C. albicans in vitro* and *in vivo*. **(A, B)** Caco2 cells were grown to confluence and layered with *C. albicans* (Hyphae-GFP) and exposed to either control vehicle or PYY 1-36 (20um) showed a significant reduction of hyphae at 6 hours. **(C)** Exogenously administered PYY_1-36_ through oral gavage significantly reduced overall *C. albicans* load in wild-type C57BL/6 mice starting at day 6 days post *C. Candida* challenge compared with scrambled peptide administered controls. (N=8/group, ANOVA (*p<0.05)). (D) PYY KO mice show increased fungal colonization of *C. albicans* compared with WT mice following *C. albicans* challenge. (N=5/group). **(E)** Mycobiome changes in the lumen and mucus layer of PYY KO vs WT animals. Beta diversity assessed by Jaccard dissimilarity displayed as principal coordinates analysis. Significance determined by permutational multivariate analysis of variance (PERMANOVA), R_2_ = 0.17, P = 0.0006, and permutational analysis of multivariate dispersions (PERMDISP2) P = 0.022. **(F)** Spearman heatmap of family and genus relative abundances of fungal taxa between sites. Dendrograms indicate experimental group and taxa clustering. **(G)** Significantly altered genus between experimental groups determined by ANOVA (*P<0.05, **P<0.01, ***P<0.005). **(H)** Alpha diversity of the fungal populations determined by Chao1 (ANOVA, P = 0.12).

Given the evidence that PC-PYY localizes to the mucus layer and may specifically modulate fungal colonization within this compartment, we next isolated luminal and mucosal associated samples from PYY-KO and WT animals to sequence the internal transcribed spacer (ITS) region. Compared with luminal fungal communities that likely carry more environmentally ingested fungi, the mucus associated layer harbored a distinct fungal population as assessed by Jaccard dissimilarity on principal coordinate analysis (Figure 6E). Within the mucus, PYY-KO animals displayed elevated colonization of *Candida* and its family, *Saccharomycetaceae*, along with *Saccharomycetaceae*, compared with WT animals (Figures 7F-H). In contrast, 16S sequencing indicated little genotype differences in mucosal associated bacterial communities in relative abundance, beta diversity, or alpha diversity (Figure S8). These data suggest the mucosal associated fungal population differs from luminal digesta and PYY may influence membership in a manner consistent with prevention of Candida colonization.

### Clinical relevance of PC-PYY to patients with Crohn’s Disease involving the terminal ileum

Several lines of investigation have supported the notion that the subset of patients with Crohn’s disease involving the terminal ileum (iCD) may have defects in Paneth Cell function (Adolph et al., 2013; Wehkamp et al., 2005). Major gene variants associated with increased risk for Inflammatory Bowel Disease (IBD) in a patients of European ancestry, for instance, have been linked to autophagic defects in these cells, and histopathological features of ileal Paneth Cells associated with iCD have been described (Cadwell et al., 2009; Stappenbeck et al., 2011; Wang et al., 2018).To explore this possibility, we performed a limited survey of iCD patients and normal controls who were sampled endoscopically. As most of the iCD patients had stricturing or severe edema of the terminal ileum that did not allow passage of the endoscope and sampling at sites of active inflammation, samples were primarily collected from sites downstream of active disease.

We found increased total fungal and *C. albicans* CFU counts in the terminal ileal mucosa of 11 iCD patients compared to 5 non-IBD controls subjects (Figure 7A). While this is a small sampling, it confirms findings from several other previously published findings, (Li et al., 2014; Hoarau, et al., 2016; Sokol et al., 2017a). Intestinal 2-dimensional (2D) epithelial organoid monolayers (In et al., 2016) were derived from stem cells of human distal ileal pinch biopsies obtained by colonoscopy in 3 iCD and 3 non-IBD control subjects. As shown in Figures 8B and 8C, these monolayers were next exposed to *C. albicans* yeast to examine whether they would adhere to the monolayers and to what extent they might transform to the more virulent hyphal forms. The quantitation of relative hyphal form abundance to total yeast in each field of view at 20 hrs is presented (Figure 7B). Filamentous *C. albicans* hyphae are clearly evident in monolayers from iCD patients in contrast to healthy cell monolayers (Figure 7C). While there are limitations to the interpretation of these *in vitro* data where cells have been passed over numerous generations in the presences of growth factors and then differentiation buffer, these findings suggest that iCD cells may be inherently permissive for fungal virulence. Our findings that PC-PYY may be important for maintenance of fungal commensalism provides a potential mechanism for this difference in permissiveness between organoids derived from healthy and control patients. Further investigations will be required to validate and extend these observations in larger human based studies. Finally, we propose a working model for PYY regulation of fungal pathogens in iCD (Graphical Abstract).

**Figure 8.**
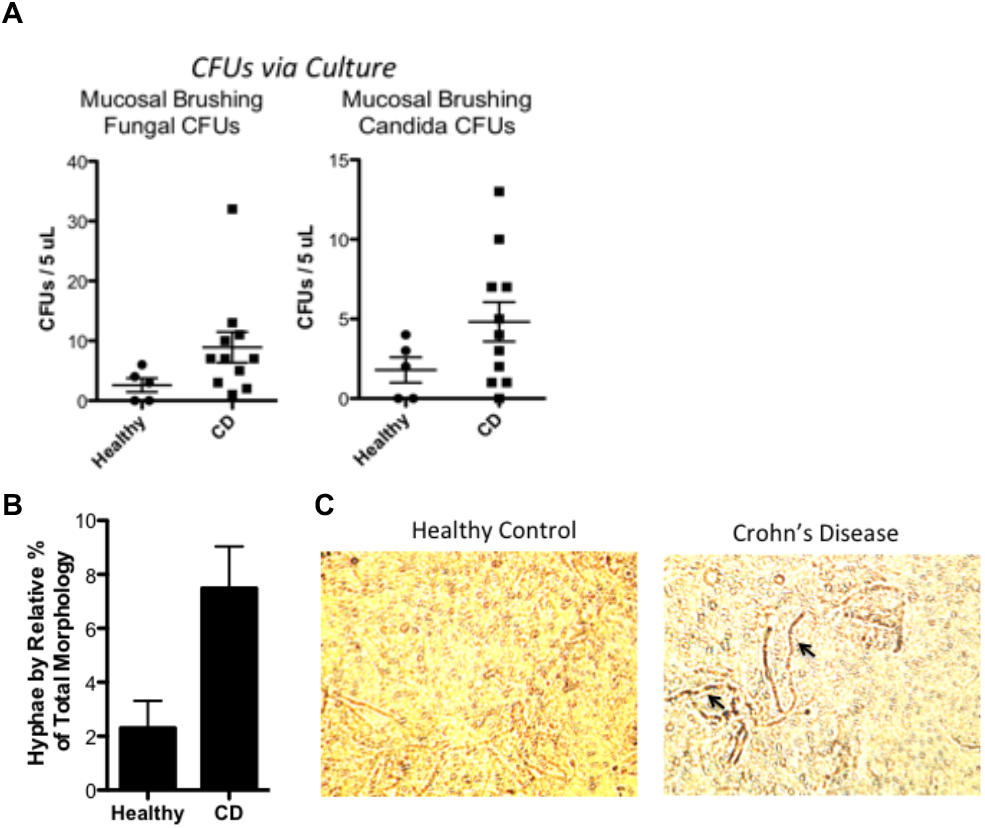
Hyphal form of *C. albicans* increased in iCD patients. **(A)** Ileal brushings and biopsies of human subjects exhibit increased fungal and *C. albicans* abundance in iCD compared with healthy controls. (B) *C. albicans* in the hyphal state adherent to mucosal surface of confluent enteroid monolayers from Crohn’s disease (CD) or healthy patients on Transwell supports coated with collagen IV. Monolayers were visualized after 20 hours of coculture with C. albicans yeasts at 37 °C. (N=3 patients/group) (C) Wells were imaged in triplicate and brightfield images of adherent virulent (hyphae) Candida (arrows) on mucosal surface of monolayers were quantified.

## DISCUSSION

Little is known about the host-mycobiome interactions in the gut. Fungi are present in the “healthy” gut microbiome, particularly the yeast *Candida albicans*, which is found in at least 70% of healthy humans (Odds, 1988). In addition to its well-known role as an opportunistic pathogen, recent studies indicate *C. albicans* may exert positive effects on immune development in healthy hosts (Jian et al., 2017; Shao et al., 2019; Bacher et al., 2019). Moreover, recent genetic analysis of C. albicans in a murine colonization model suggests that the host actively curate the fungal population by restricting *C. albicans* growth in response to excess antigenic markers of invasive hyphae (Witchley et al., 2019). In this setting, our discovery of unanticipated antimicrobial activity of the well-known satiety hormone, PYY, seems particularly relevant. Peptide YY (PYY) is a 36 amino acid-residue peptide that is highly conserved in vertebrate species, where its structure is virtually unchanged between cartilaginous fish and humans (Conlon, 2002). Since its discovery in porcine intestine and brain stem tissue in 1982 (Glavas et al., 2008; Tatemoto, 1982), diverse regulatory roles have been identified for PYY, mostly involving metabolic regulation (Batterham et al., 2007; Hill et al., 2011; Holzer et al., 2012). In the mammalian and human intestine, PYY is notably released into the bloodstream by ‘L cells’, a subset of enteroendocrine cells (EECs) that are most numerous in the distal ileum and colon (Gunawardene et al., 2011). PYY_3-36_ concentrations in circulation increase after meals and decrease with fasting, supporting its function as a satiety peptide for regulating hunger and mediating gut physiology (Murphy and Bloom, 2006). Here, we report that PYY is also expressed in distinct, non-lysosomal secretory granules of murine and human Paneth cells. Unlike L cells, Paneth cells play important roles in innate immunity by producing antimicrobial peptides that regulate microbial assemblage and deter pathogens. The presence of PYY in PC secretory granules therefore raised the possibility that PYY has a previously unrecognized biological function, that of an antimicrobial peptide. PYY also has structural features and charge characteristics that are strikingly similar to the amphibian antimicrobial peptide, magainin-2. We therefore set out to determine if PYY had antibacterial activity. Surprisingly, PYY has modest effects and failed to inhibit the proliferation or viability of a variety of bacteria even at high doses. Nevertheless, based on published observations that magainin-2 on the skin of aquatic frogs has anti-fungal properties, we examined the *in vitro* activity of PYY against fungi. As described above, *Candida albicans* is the primary fungus associated with the human gut. *C. albicans* can adopt multiple cell morphologies, with round-to-oval, single celled yeasts behaving as commensals and highly elongated, multicellular hyphae and pseudohyphae associated with virulence in the GI milieu (Sudbery et al., 2004; Staib and Morschhäuser, 2007; Sudbery, 2011; Thompson et al., 2011). Interestingly, the addition of either PYY has specific activity against the hyphal (virulent), but not yeast (commensal), forms of *C. albicans* under hyphae-inducing conditions. PYY also inhibits cell respiration in hyphae, but not yeast, and inhibits biofilm formation. The mechanism of fungal cell injury by PYY appears to involve direct membrane disruption mediated by charge interactions at the surface, similar to other AMPs, as uptake of propidium iodide occurs where FITC-labeled PYY binds the hyphal membranes and causes membrane blebbing and irregularities. Moreover, the anionic surface charge of C. albicans hyphae, in contrast to the absence of this in the yeast form (Figure 4A) attracts PC-PYY binding through electrostatic interactions. From these data, we conclude that: (1) PC-PYY differs from the endocrine-PYY in structure, function, action, (2) PC-PYY is a novel and functionally specific AMP that specifically targets the virulent form of *C. albicans* to maintain a healthy state of fungal commensalism in the gut, (3) PC-PYY is packaged in secretory granules that are distinct from those that carry lysozyme and is released by mucosal exposure to intact hyphae, and not yeast *C. albicans*, (4) the specificity of the cationic PC-PYY for the former lies in its initial electrostatic interaction with the anionic surface charge of *C. albicans* hyphae, which the yeast form does not have, (5) once secreted PC-PYY is retained by overlying mucus for which it has proclivity and which in turn provides optimal conditions for its activity and protection from DPP-IV cleavage, and (6) Mucus fortified with PC-PYY kills and prevents further penetration by virulent *C. albicans*. At the same time, PC-PYY has limited effects on commensal *C. albicans* yeast and other members of the gut microbiome. This microbial selectivity, activation, and mucus compartmentalization of PC-PYY distinguishes it from other innate immune AMPs which have broad ranges of action (Bevins and Salzman, 2011; Mukherjee and Hooper, 2015), representing an elegant mechanism by which the host can prevent the development of pathogenesis while retaining its commensal fungal mycobiome.

The dual functions of peptide YY as an innate antimicrobial peptide and an anti-satiety hormonal peptide bears special mention. PYY joins a growing list of mammalian proteins and peptides that have been demonstrated to serve multiple discrete functions within the host (Broderick, 2015). In the case of PYY, this involves a proteolytic (DPP-IV) cleavage of the two N-terminal amino acids, transforming the AMP form, PYY_1-36_, that utilizes biophysic (electrostatic and lipid disruption) action to kill *C. albicans* hyphae to the hormone form, PYY_3-36_, that binds and signals through a Y2-membrane receptor (Batterham et al., 2002, 2007; Holzer et al., 2012). In this regard, there is little cross-over in function by these two PYY entities. The major circulating form, PYY_3-36_, has limited anti-fungal activity, whereas, PYY_1-36_, has far less binding affinity for the Y2 receptor and hormonal activity (Batterham et al., 2006). The repurposing of the PYY molecule appears dependent on non-genomic factors that include the nature of extracellular stimulus, systemic vs luminal-directed secretion, the milieu (mucus vs plasma), and mucus conferred protection of the PYY_1-36_ from enzymatic cleavage by DPP-IV which would convert it to an endocrine form.

Finally, we believe our findings may have relevance to human health and disease. In health, expression of PC-PYY in humans has been corroborated which likely serves as a unique antimicrobial peptide that plays a role in maintaining a commensal state in a critical transitional zone between vastly different small and large bowel microbiota. Weakening or dysfunction of Paneth cells in the terminal ileum (and appendix) can perturb the often delicate balance between host and microbe, resulting in increased risk or frank development of disease, particularly in genetically susceptible hosts. In IBD, particularly Crohn’s disease involving the terminal ileum, Paneth cell dysfunction and fungi have been implicated in disease pathogenesis, but the supporting evidence is minimal and often circumstantial. Several studies have reported increased titers of *C. albicans* in stool samples of ileal CD patients (Sokol et al., 2017a). Serum anti-fungal antibodies, particularly ASCA (anti-*Saccharomyces cerevisiae* antibody), also appear to be predictive of risk and severity of CD (Israeli et al., 2005; Ksiadzyna et al., 2009). In fact, ASCA is the single most robust biomarker manifestation of CD (Barnes et al., 1990; McKenzie et al., 1990; Giaffer et al., 1992; Quinton et al., 1998),particularly involving gastroduodenal and small bowel rather than colonic disease and associated with the likelihood of having more severe disease requiring surgery within a 9-year follow-up period (Walker et al., 2004). ASCA is directed against mannan, a polysaccharide that decorates the cell walls of many fungi, including *C. albicans*. The induction of IgG against this cell wall component is thought to reflect fungal invasion of the gut epithelium, a property not associated with normal commensalism. In addition, mice lacking Dectin-1, an innate immune pattern-recognition receptor for another fungal cell wall component, ß-1,3-glucan, have increased susceptibility to chemically induced colitis, attributed to increased epithelial invasion by Candida species. Human polymorphisms of Dectin-1 (CLEC7A) and CARD9, both part of an innate immune signaling pathway, are strongly linked to a severe form of ulcerative colitis, autoimmune disorders, and Candidiasis (Khor et al., 2011; Jia et al., 2014; Underhill and Iliev, 2014; Lanternier et al., 2015). Despite these intriguing hints of a role for fungi in IBD, the behavior of fungi within the gut has been largely uncharacterized.

Crohn’s disease of the terminal ileum (iCD) has also been associated with Paneth cell dysfunction/injury and histopathology (Liu et al., 2016). In some, but not all, these defects have been linked to IBD gene risk variants associated with aberrations in autophagic functions (e.g. NOD2, ATG16L1 T300A, IRGM, and LRRK2 (Anderson et al., 2011; Fritz et al., 2011; Khor et al., 2011; Van Limbergen et al., 2014; Lanternier et al., 2015). Defective Paneth cell function can lead to the development of intestinal dysbiosis (Wang et al., 2018; Wehkamp et al., 2005). Both Paneth cell dysfunction and mucus-depletion (which is common in chronic inflammation of the gut) could result in decreased PC-PYY activity, permitting common fungal commensals like *C. albicans* to transform into virulent hyphae that can attach, form biofilm, invade the mucosal barrier and mucosa, and trigger or contribute to the development of iCD. We attempted to explore this possibility using both *in vitro* and *in vivo* approaches and demonstrated (1) PYY_1-36_ inhibits *C. albicans* hyphae from attaching to Caco2 colonic epithelial monolayers, (2) increased clearance of *C. albicans* in a murine model with oral PYY administration, (3) increased *C. albicans* colonization in PYY^-/-^-gene deficient mice, (4) increased *C. albicans* load from ileal samples acquired endoscopically from patients with active iCD, and (5) increased attachment of *C. albicans* hyphae to 2D-organoid monolayers from iCD patients compared to non-IBD controls 2D monolayers. Unfortunately, direct measurements of mucus and mucosal PYY levels were technically not possible because the narrow passage caused by edema and fibrosis did not permit intubation of the endoscope into active areas. Collectively, these data suggest that iCD patients have an increased *C. albicans* load and that PYY, both *in vitro* and *in vivo*, can deter *C. albicans* attachment to the mucosa.

In summary, we propose that PC-PYY is a unique antimicrobial peptide that differs from its endocrine counterpart in structure, biological action, and function. While it is found in virtually all vertebrate lineages, these observations raise questions about the origins of endocrine molecules and about nature’s repurposing of conserved molecules for multiple homeostatic purposes in the host. Regulation and selective actions of PYY to the virulent form of *C. albicans* further distinguishes it amongst other innate immune peptides, implicating a role in regulating key elements of the gut mycobiome. In active iCD, it may very well be that PC-PYY bioavailability, stability, and bioactivity are compromised by Paneth cell dysfunction and mucus cell depletion, providing fungi like *Candida albicans* with the opportunity to become virulent, adherent, and invasive, and to contribute to the etiopathogenesis of the disease.

## ACKNOWLEDGMENTS

We acknowledge the MRSEC Shared User Facilities at the University of Chicago (NSF DMR-1420709) and the Mass Spectrometry Facility (NSF CHE-1048528). The authors thank Daina L. Ringus, Hyoann Choi, Mirae Lee and Monika Krezalek for helpful discussions. We would like to acknowledge Vytas Bindokas at the Integrated Light Microscopy Core. We thank the Garvan Institute of Medical Research and Dr. Herzog for generously providing the PYY knockout line (Batterham et al., 2006)

## AUTHOR CONTRIBUTIONS

JFP and EBC were involved in every aspect of this study, including the discovery, conception, experimentation, data analysis, and MS preparation; SMN, JW, VL, HH, DLT, AS, KGH, AK, XZ, YT, CMC, OZ, JA, and LW each had a key role in contributing to the conceptual, experimental, and data analysis components of this study.

## Supplemental Figures

**Figure S1.**
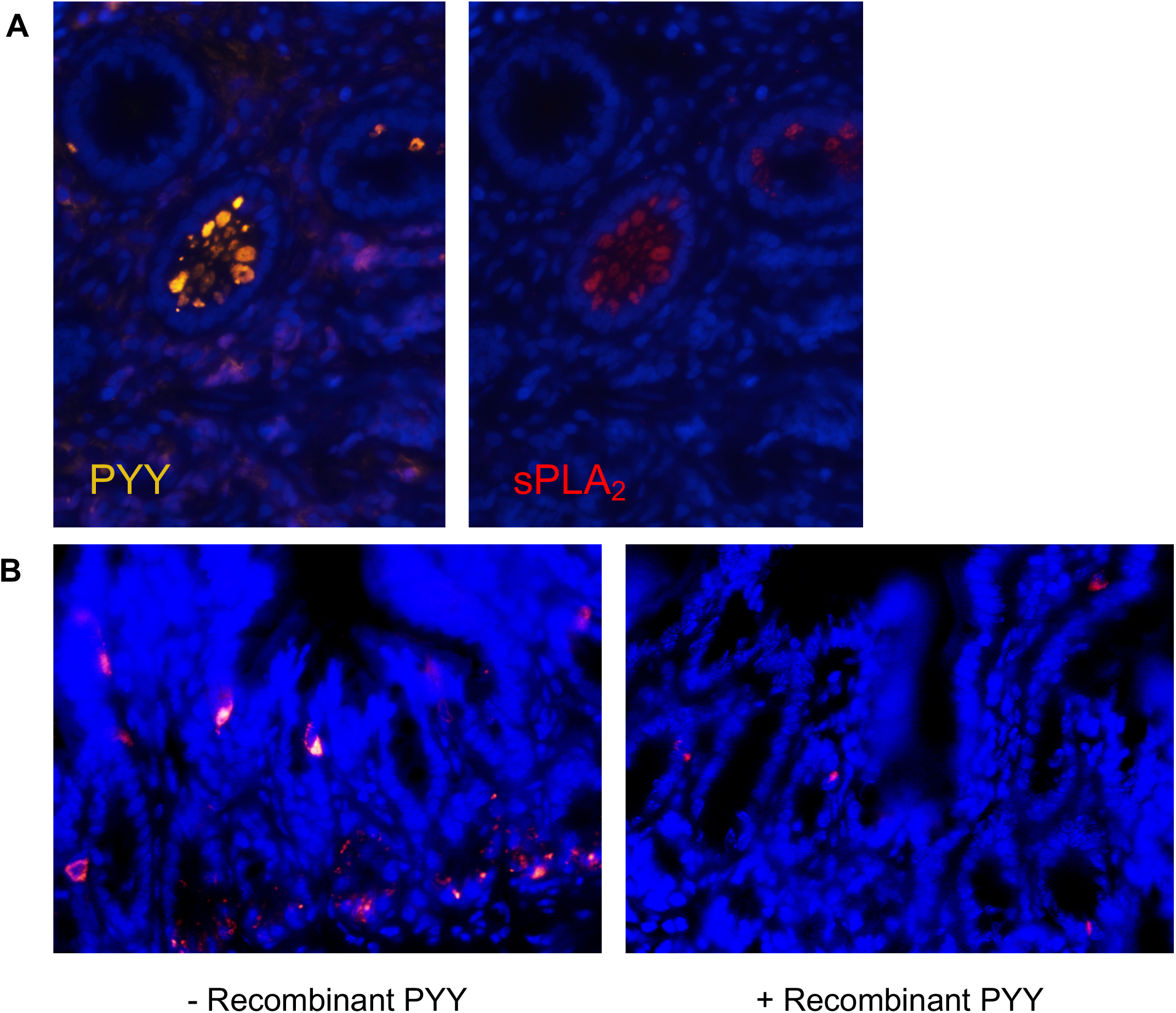
PYY immunolocalization to Paneth cells. **(A)** PYY (yellow) immunolocalized in human ileal biopsies to the human Paneth cell with secretory phospholipase A_2_ (red, sPLA_2_). (B) Primary PYY antibody pre-adsorbed with recombinant PYY results in near complete quenching of epitope recognition and signal from both L-and Paneth cells.

**Figure S2.**
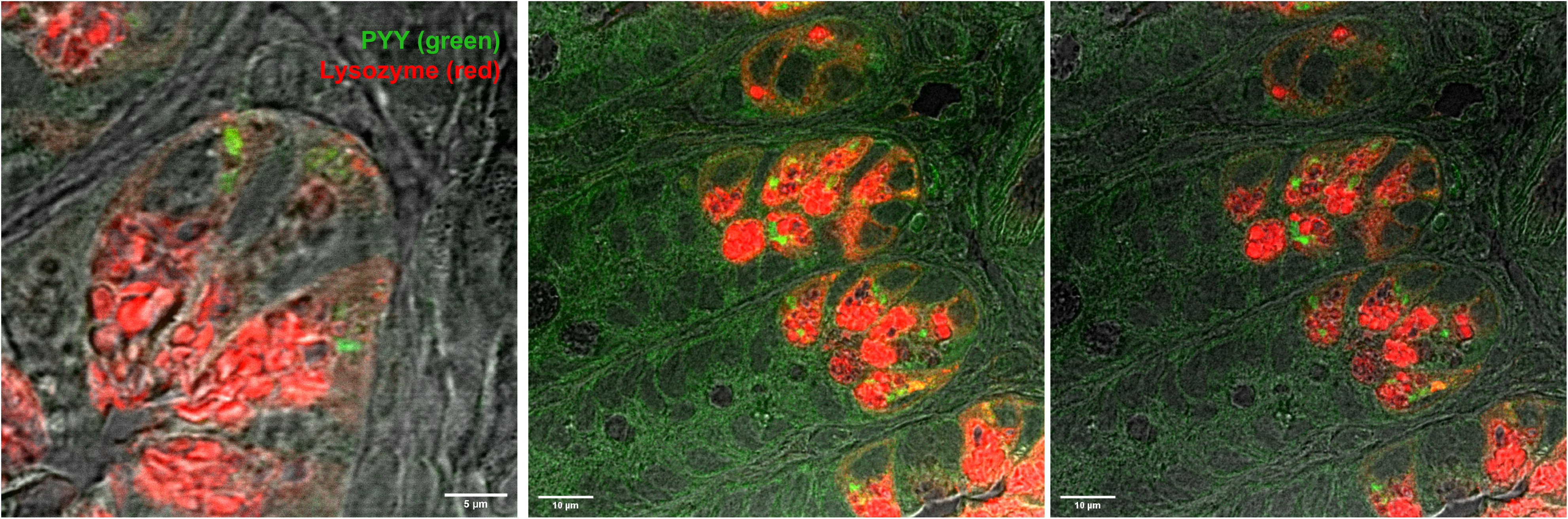
PYY and lysozyme localization. PYY (green) and lysozyme (red) do not colocalize and are packaged in separate secretory granules of ileal Paneth cells within the small intestine. Images acquired using STED Sp8 confocal microscopy.

**Figure S3.**
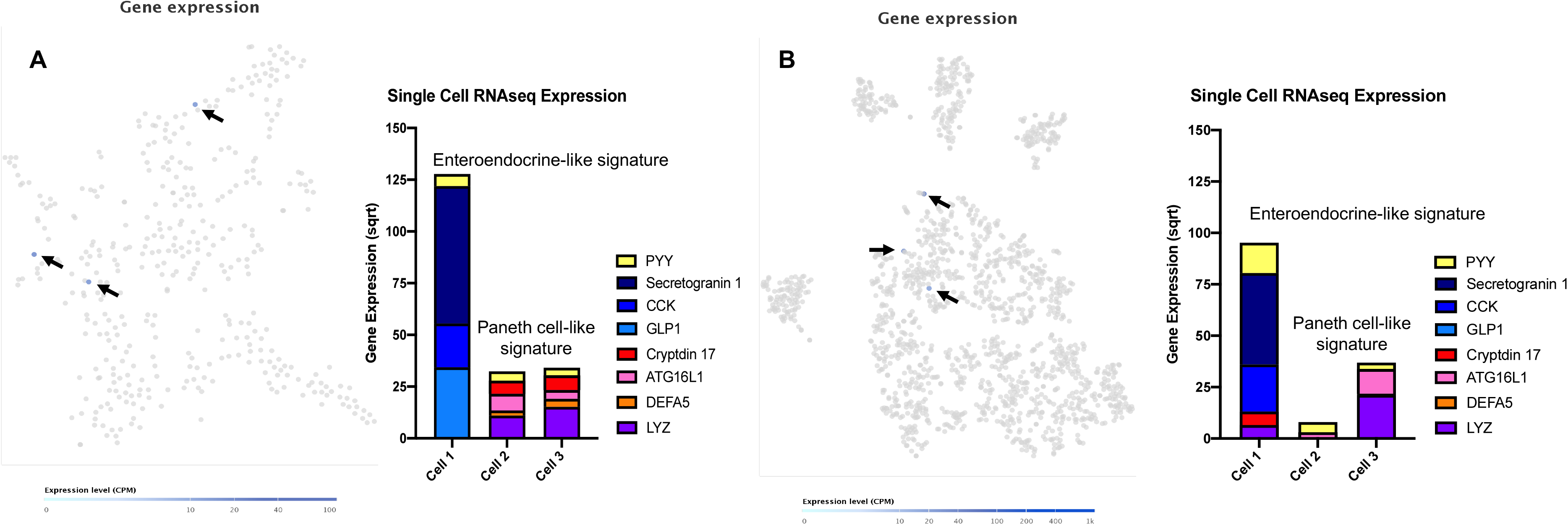
Single cell RNAseq from murine small intestine. Single cell RNAseq data from murine small intestinal epithelial cells was utilized to characterize PYY expressing cells. **(A)** 387 cells and **(B)** 1522 epithelial cells from publicly available databases (Haber et al, Nature 2017) were first searched for PYY expression (black arrows). The co-expression of enteroendocrine products (Secretogranin-1, CCK, and GLP1) and Paneth cell products (Cryptdin 17, ATG16L1, DEFA5, and Lysozyme 1) were also determined. PYY expressing cells were characterized by either an Enteroendocrine or Paneth cell signature based on co-expression of key marker genes.

**Figure S4.**
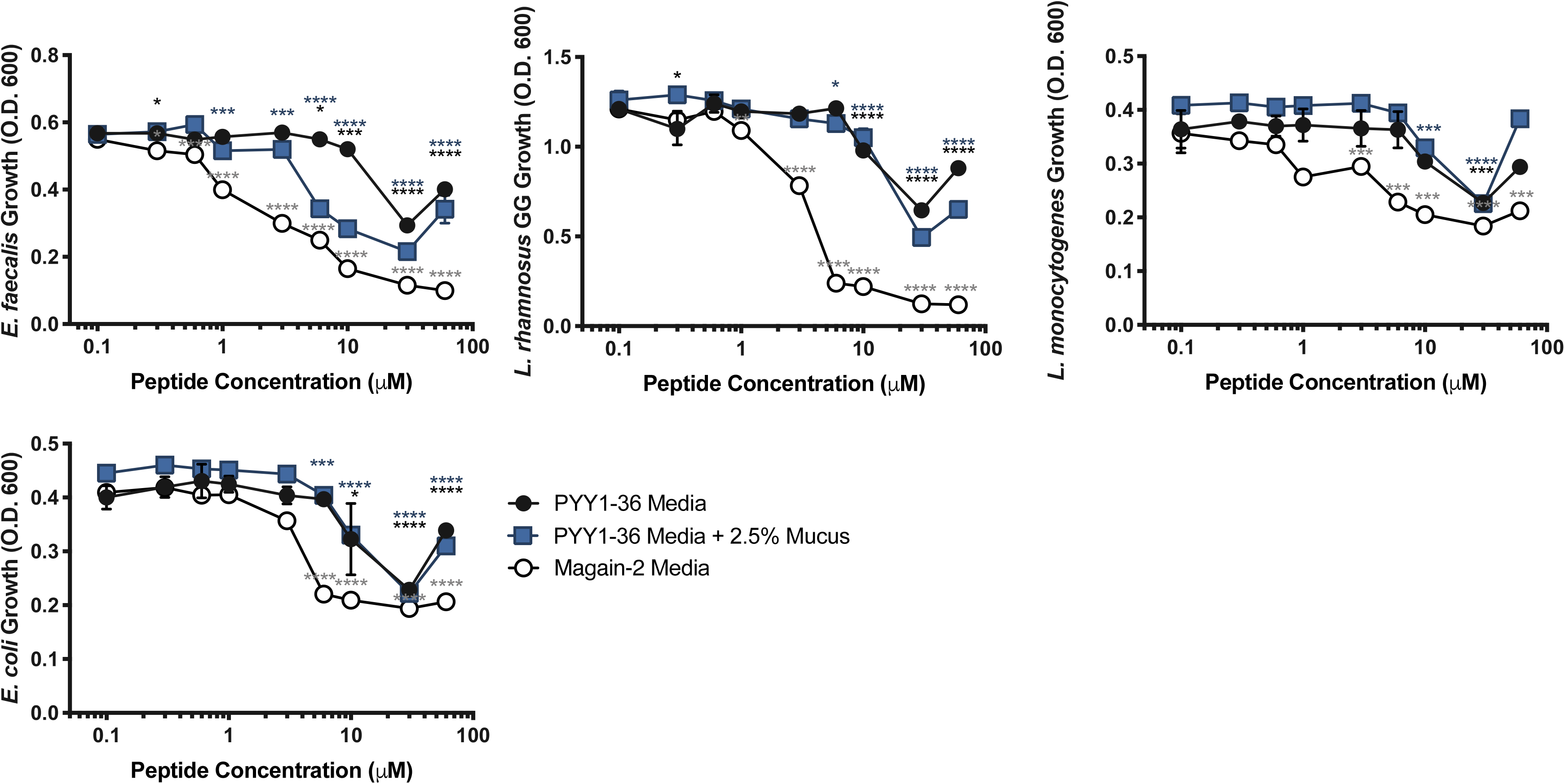
PYY 1-36 inhibits growth of certain bacterial strains. Exposure to varying concentrations of PYY_1-36_ in specific bacterial growth media and media with 2.5% w/v porcine stomach mucus impacts growth of certain bacteria. Effects were biphasic with the greatest inhibition around 30 μM. All assays performed in duplicate three times, O.D. values were normalized to the plate containing 0 μM PYY and significance was analyzed by ANOVA (*p<0.05; **p<0.01; ***p<0.001; ****p<0.0001).

**Figure S5.**
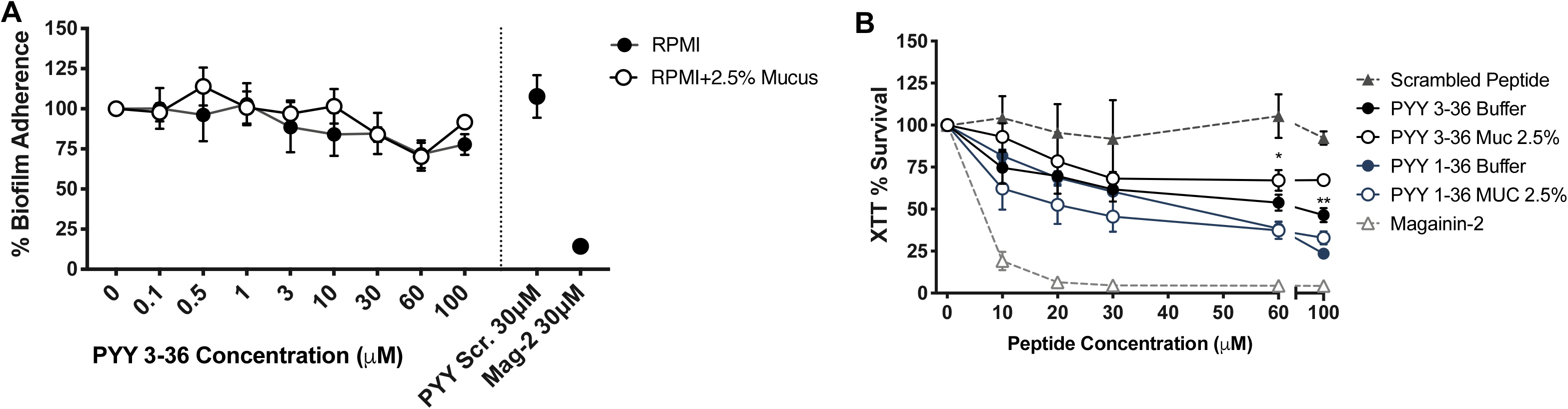
PYY_3-36_ Impact on percent *C. albicans* biofilm adherence and survival. **(A)** No significant decrease in percent biofilm adherence was observed when *C. albicans* was treated with PYY_3-36_ in either RPMI or mucus compared to magainin-2. (B) PYY_3-36_ in buffer (black circle) and mucus (black open circle) showed less impact on hyphae viability compared to PYY_1-36_ in buffer (navy circle) and mucus (navy open circle). All assays performed in triplicate two times, O.D. values were normalized to the plate containing 0 μM PYY and significance was analyzed by ANOVA (*p<0.05; **p<0.01; ***p<0.001; ****p<0.0001).

**Figure S6.**
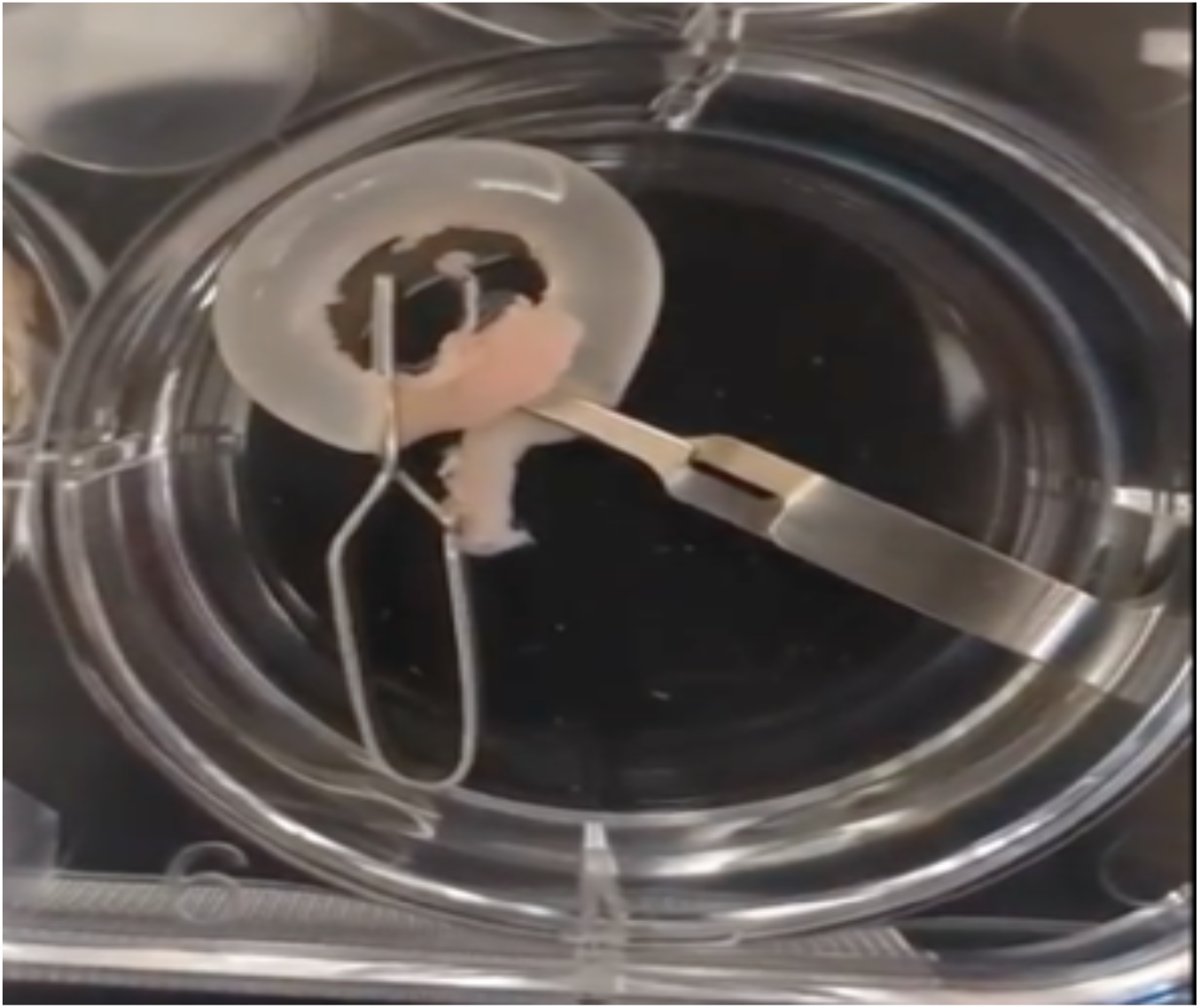
Preparation of ex *vivo* distal ileum loop. Distal ileal sections were dissected from the mesentery, rinsed, and ligated on one end. Loops were filled with stimulant, ligated on the other end, and then placed in a well with buffer. After 2 hours of incubation, tissues were immediately processed for analysis (well, lumen, mucus, tissue).

**Figure S7.**
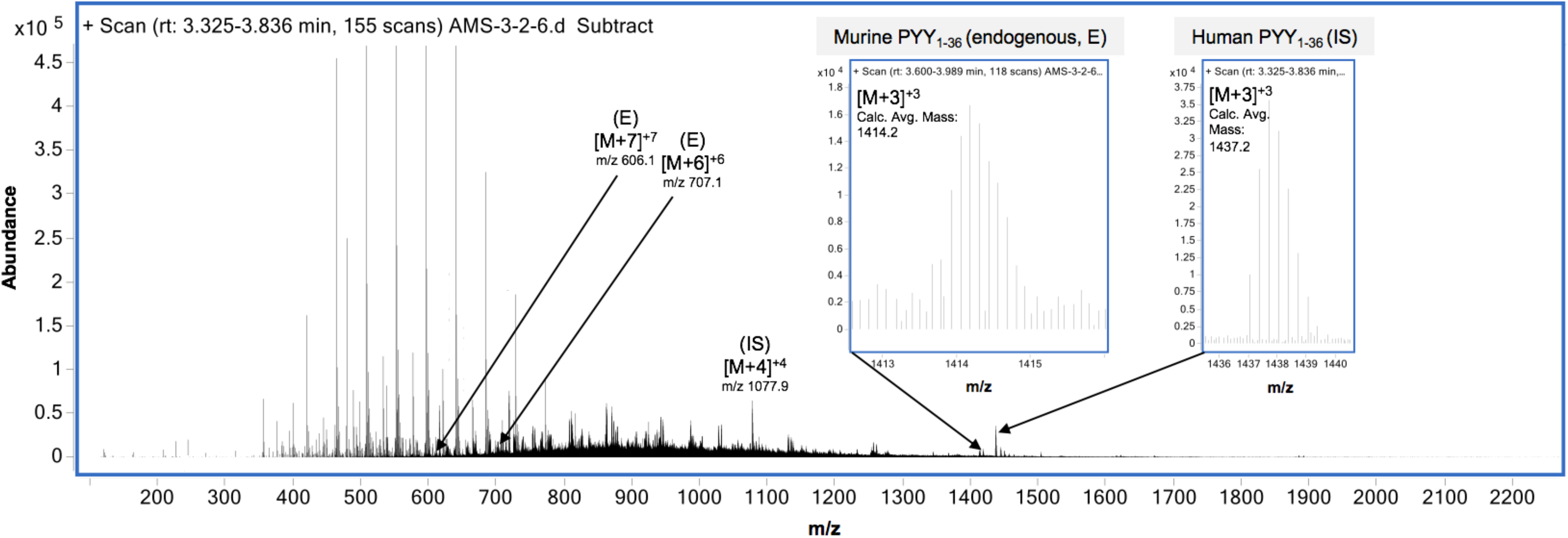
Spectra of murine PYY_1-36_ and human PYY_1-36_ ions quantified in ex *vivo* ileal loop samples by UPLC-ESI-QTOF-MS. Human PYY_1-36_ m/z 1077.9 [M+4]^+4^, 1437.2 [M+3]^+3^; Human PYY_3-36_ m/z 1350.5 [M+3]^+3^; Murine PYY_1-36_ m/z 1414.2 [M+3]^+3^, 606.1 [M+7]^+7^; Murine PYY.

**Figure S8.**
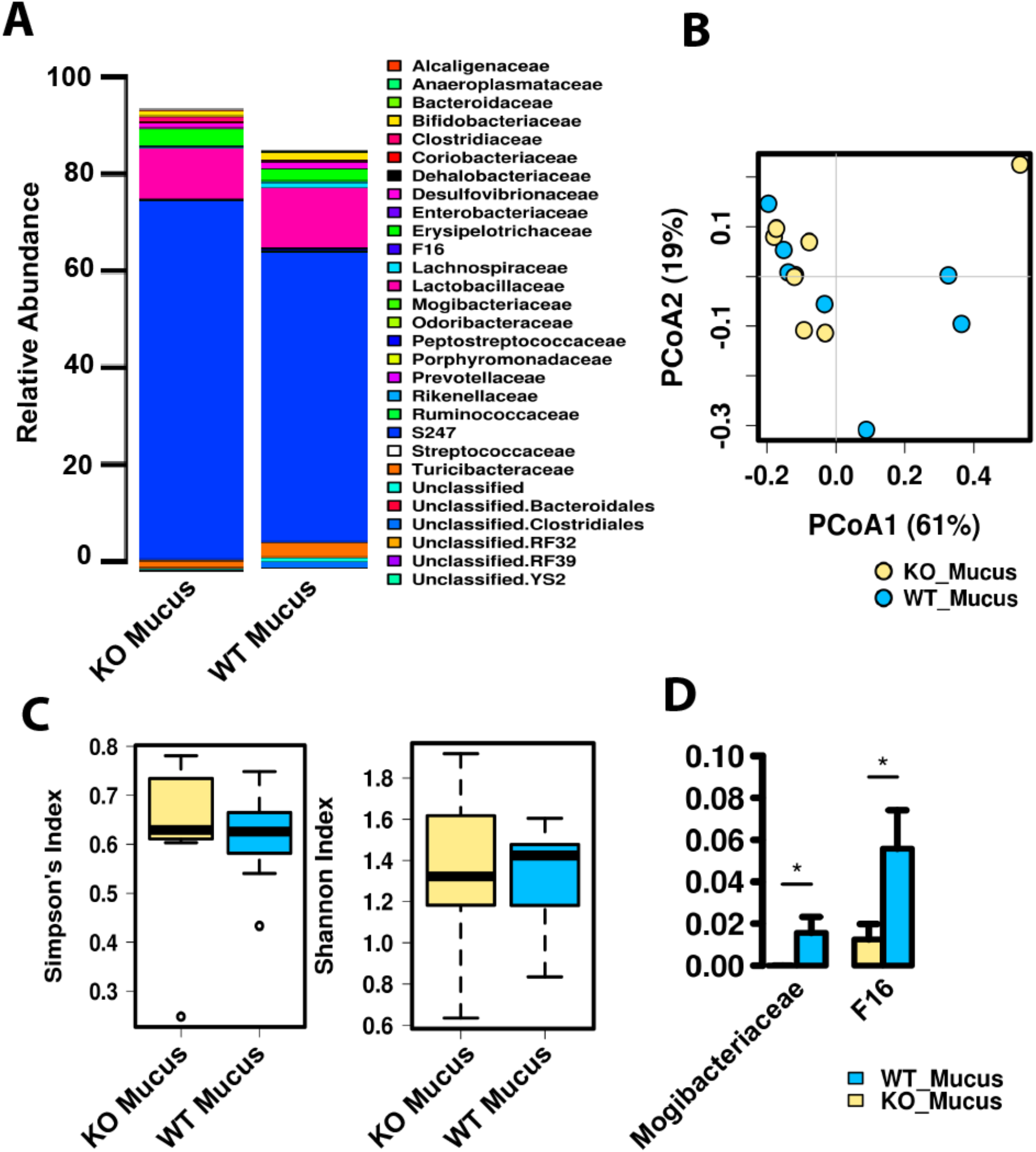
Microbiome of mucous layer between PYY KO vs WT animals. **(A)** Relative abundance of family level taxa between groups. **(B)** Beta diversity assessed by Bray-Curtis dissimilarity displayed as principal coordinates analyses. Significance determined by permutational multivariate analysis of variance (PERMANOVA), R_2_ = 0.034, P = 0.78, and permutational analysis of multivariate dispersions (PERMDISP2), P = 0.55. **(C)** Alpha diversity of the fungal populations determined by Simpson’s (P = 0.91) and Shannon Indexes (P = 0.84). **(D)** Significantly altered taxa at the family level between KO and WT animals. (*P<0.05).

