## Supplemental Materials and Methods for "Peptide YY: a novel Paneth cell antimicrobial peptide that maintains fungal commensalism"

**STAR METHODS**

**RESOURCE AVAILABILITY**

**Lead Contact**

The lead contact for our study is Dr. Eugene Chang. For further information and requests of resources please contact him at.

**Materials Availability**

This study did not generate new unique reagents or mouse lines.

**Data and Code Availability**

The ITS and 16s datasets generated during this study were deposited at the National Centre for Biotechnology Information Sequence Read Archive (accession number SUB7307311).

For the single cell RNAseq analysis, datasets from (Haber et al., 2017) were utilized.

**EXPERIMENTAL MODEL AND SUBJECT DETAILS**

**Animals**

All animal protocols were approved by IACUC at the University of Chicago. Animals were either wild-type (WT) or PYY^-/-^ gene-deficient mice on the C57Bl/6 background bred and housed under standard 12:12 light/dark conditions at the University of Chicago (Shi, et al., 2015)(Batterham et al., 2006). In *Candida* challenge experiments, male C57BL6/J mice aged 7-9 weeks old, weighing 18-22 grams were used and maintained on Harland Teklad feed. Furthermore, studies using *Candida spp*, animals were maintained in single BSL2 cages. Germ-free animals were maintained in sterile isolators in our gnotobiotic facility.

**Human Ileal Biopsies**

Human ileal pinch biopsies were collected from patients with Crohn’s Disease undergoing colonoscopy and fixed in 4% formalin/PBS or used to generate enteroids. Procedures were performed under IRB approval at the University of Chicago (IRB #15573A) and with written consent from all patients. Biopsies were used to generate intestinal organoid 2D monolayers, as described (In et al., 2016). Following confluence of monolayers, 40 μL (1.0 OD_600_) of *Candida albicans* (HGFP3, GFP tagged HWP1, Gift from UChicago Dr. John Alverdy) from overnight cultures were washed, resuspended, and inoculated per 2 mL well and cultured for 20 hours at 37°C in enteroid growth media. Wells were imaged in triplicate fields and the presence of hyphae was confirmed under GFP fluorescence.

**Microbial Strain Culture**

*Lactobacillus rhamnosus* GG (ATCC 53103), *Enterococcus faecalis (*ATCC BAA-2128), *Listeria monocytogenes* EGD (ATCC 49594), *Peptostreptococcus* *anaerobius* (ATCC 27337), *Staphylococcus aureus* Newman *(*ATCC 25904), *Escherichia coli* K12 (ATCC PTA-7555), *Bacteroides fragilis* (ATCC 25285), *Salmonella enterica (*ATCC 1575D-5) and *Pseudomonas aeruginosa* MPAO1-P2 (Luong et al., 2014) gift from Dr. John Alverdy, University of Chicago) were streaked out from frozen glycerol stocks onto either brain heart infusion broth (BHI, *E. faecalis*, *L. monocytogenes*, *S. aureus*: BHI - Fisher Scientific, Cat# 237500), brain heart infusion-supplemented (BHIS, *P. anaerobius, B. fragilis*: BHIS - ATCC Medium 1293), Luria broth (LB, *E. coli* K12, *S. enterica*, *P. aeruginosa*: LB - BD Difco, Cat# DF0446-07-5), or De Man, Rogosa and Sharpe broth (MRS, *L. rhamnosus* GG: MRS - BD Difco, Cat# DF0881175). Single colonies were inoculated into their respective liquid media and cultivated under aerobic and anaerobic conditions at 37°C for 24 – 48 hours.

**Fungal Cultures**

*Candida albicans* SC5314 (ATCC MYA-2876) was maintained in frozen glycerol stocks at -80°C and streaked out on yeast-peptone-dextrose (YPD, Sigma-Aldrich, Cat# Y1375) agar plates for 48 hours at 30°C prior to use. Single colonies were selected and cultured in YPD media at 30°C overnight with gentle shaking. To induce hyphal growth, overnight yeast cultures were placed into RPMI 1640 Medium (Thermo Fisher Scientific, Cat# 21870-076) (1:100) at 37°C for 4 hours. For yeast, overnight yeast cultures were diluted (1:100) in YPD or RPMI and kept at room temperature to prevent hyphal transformation. *C. albicans* HGFP3, the strain in which gene encoding GFP was inserted after the promoter of HWP1 in the SC5314 strain. Since HWP1 is known to be expressed by hyphae, the *C. albicans* HGFP3 will express GFP only when the cells grow as true hyphae (Chairatana et al., 2017).

**METHOD DETAILS**

**Murine *Candida albicans* Challenge**

Mice were administered clindamycin (PO gavage, 25 mg) and cefoxitin (IM, 50 mg) 2 days before oral *Candida* challenge. One day prior to challenge, mice were fasted and administered clindamycin (25 mg). Mice were gavaged with 200 µL of 1.0 OD_600_ *Candida albicans* (SC5314) in the morning and provided food 4 hours later. Clindamycin (25 mg/day) was administered on days 0, 2, 4, 6, and 8 until sacrifice. In studies examining oral peptide administration, peptides (200 µg) were suspended in sterile PBS (100 µL) and gavaged on days 2, 3, 4, and 8. Fecal pellets were collected on days -2, 0, 2, 4, 6, and 8, weighed, resuspended in sterile PBS and plated in duplicate on YPD plates (Sigma-Aldrich, Y1375) containing gentamicin (0.1 mg/mL) and vancomycin (0.01 mg/mL), to prevent bacterial growth, at 30^o^C. Growth of *Candida spps* was confirmed on CHROmager *Candida* selection plates (BD, Spark, MD, Cat# 8012620). Animals were humanely euthanized, and regions of small and large bowel tissues were taken for analysis.

**Immunofluorescence and Epitope Blockade**

Mouse ileal tissues were fixed in 4% formalin/PBS or, to preserve mucosal area, in methyl Carnoy’s solution (60% ethanol, 30% chloroform, and 10% glacial acetic acid) overnight at 4°C. Tissues were then embedded in paraffin. Human pinch biopsies were fixed in 4% formalin/PBS and embedded in paraffin. Five-micron sections were cut, deparaffinized, rehydrated, and washed in PBS. Antigen retrieval was performed by boiling sections in sodium citrate (pH 6.0) for 10 mins and cooling to room temperature. Tissue sections were blocked with Protein Block (Agilent, DAKO, Cat# X090930-2) for 45 mins followed by anti-PYY antibody (1:200, rabbit polyclonal anti-PYY, Cat# ab22663, Abcam) and anti-lysozyme antibody (1:400, goat anti-lysozyme, Santa Cruz Biotechnology, Cat# C-19) in mice or anti-PYY (1:200, rabbit anti-PYY, Abcam, Cat# ab22663) and anti-SPLA2 (1:250, mouse anti-sPLA2 group 2, Santa Cruz, Cat# SCACC353) in human sections and incubated overnight at 4°C. Samples were washed and incubated with conjugated secondary antibody (1:1000, Alexa Fluor 555 donkey anti-rabbit IgG, Thermo Fisher Scientific, Cat# A-31572; 1:1000, Alexa Fluor 647 donkey anti-goat IgG, Thermo Fisher Scientific, Cat# A-21447; 1:1000 Alexa Fluor 647 goat anti-mouse IgG, Thermo Fisher Scientific, Cat# A-21240) for 1 hour at room temperature. For dipeptidyl peptidase-IV (DPP-IV) detection, sections were labeled with primary goat anti-DPP-IV/CD26 antibody (1:250, RandD Systems, Cat# #AF954) and conjugated secondary antibody (1:1000, Alexa Fluor 647 donkey anti-goat, Life Technologies, Cat# A-21447). Lectin from *Ulex europaeus* agglutinin (UEA-1)-fluorescein isothiocyanate (FITC) conjugated (1:500, Sigma-Aldrich, Cat# L9006) was used to stain for mucus. Slides were counterstained with DAPI and visualized with a Leica DM2500 microscope (Leica Microsystems, Wetzlar, Germany) through a 20X lens objective using Image Pro-Plus software (Media Cybernetics, Silver Springs, MD, USA) for image capture. To confirm epitope specificity, full-length recombinant immunizing peptide was added to the primary anti-PYY antibody prior to staining followed by the remaining protocol as stated.

**Fluorescent In Situ Hybridization (FISH)**

Sections were deparaffinized and rehydrated. Antigen retrieval was performed as described above. To stabilize the tissue and nucleic acids after permeabilization, sections were re-fixed with 4% paraformaldehyde for 10 mins, washed, and dehydrated. To denature the FISH probe for PYY (IDT: 5’ cy5/ GTGATGGAGTTGGACCAGTG), it was placed in hybridization buffer (1:100, 6M NaCl, 1M Tris/HCl, 35% Formamide, 10% SDS) at 80°C for 5 min and placed immediately on the slide or on ice. Slides were incubated with the PYY FISH probe at 50°C overnight in a sealed, moist container. Sections were then stringently washed and stained for lysozyme, counterstained with DAPI, and mounted with Prolong Gold Anti-fade (Life Technologies, Thermo Fisher Scientific, Cat# P36930).

**Super Resolution Microscopy**

For super resolution microscopy, 170 μm cover slips were coated with poly-L-lysine overnight. Next, five-micron tissue sections were cut and placed on slides and allowed to dry. Deparaffinization, rehydration, antigen retrieval, blocking, and primary antibody steps were performed as described above, but with higher primary antibody concentrations; 1:100 rabbit anti-PYY and 1:100 donkey anti-lysozyme in antibody diluent (Agilent Technologies, Cat# S0809). Next, slides were washed five times (5 mins each) in PBS. Samples were fixed in 4% paraformaldehyde for 20 min at room temperature and washed in PBS three times. Secondary antibody was added for 1 hour as above and washed three times in PBS. Slides were kept in PBS and mounted immediately before imaging on a Leica Super-Resolution System (SR GSD 3D) and analyzed with ThunderSTORM software (Ovesný et al., 2014). Specifically, crypt bases were imaged to visualize individual Paneth cells containing lysozyme, and villus L-cells containing no lysozyme.

**Laser Capture Microdissection (LCM)**

To investigate whether PYY mRNA expression was present in the Paneth cell compartment, we performed laser capture microdissection (LCM). Optimal cutting temperature compound (OCT)-embedded frozen ileal samples were cut 8 μm thick and placed on frozen PEN slides (Leica Microsystems, Cat# 11505158). Slides were processed rapidly, with 30 s in Carnoy’s solution, 1 min in 70% ethanol, and 10 submersions in ddH_2_O. Sections were stained with Alcian blue for 1 min, dipped in ddH_2_O, and placed in nuclear fast red for 30 s. Slides were then quickly dipped in various ethanol solutions (70%, 90%, 95%, and 100%) and dried. LCM was performed on a Leica DM 6000B microscope. Regions of villous epithelium or the crypt base, as visualized by the presence of Paneth cell granules, were isolated and collected into separate tubes and labeled villous and crypt fractions, respectively. Total RNA was isolated using the RNeasy Micro Kit (Qiagen, Cat# 74004). RT-PCR was performed to quantify PYY and various other proteins specifically found in either the crypt (lysozyme and cryptidin-1) or villus (sucrase-isomaltase and neurotensin) of the mouse ileum (N=6, repeated twice). Primers used for RT-PCR are as follows and were purchased from Integrated DNA Technologies (IDT): PYY, Forward -GTTAACTACACCGACTTCACT: Reverse- GTCCGAGACACCGAGATA; Lysozyme (Lyz1), Forward - ATAAATTCTCAGCTCATGTGTC: Reverse- TCTTCTGTCTCAGAGTTAGTATG; Cryptdin 1, Forward AAGAGACTAAAACTGAGGAGC: Reverse GGACACCAGTACTCACATTC; Sucrase Isomaltase, Forward TATGGTGGGAATGAACAACAG: Reverse CAGGAATTCAAACTGATGGTG.

**Single Cell RNAseq gene expression**

To assess individual intestinal cell RNA gene expression, we searched the <https://www.ebi.ac.uk/gxa/sc/home> database for experiments that included adult murine intestinal epithelial cells. Two experimental subsets meeting criteria were identified, including datasets with 387 and 1522 intestinal epithelial cells from (Haber et al., 2017). PYY expressing cells were first identified and subsequently assessed for expression of Secretogranin-1, CCK, GLP1, Cryptdin 17, ATG16L1, DEFA5, and Lysozyme 1. The square root of respective gene targets was calculated and graphed.

**Peptide Modeling**

To predict the structure of peptides, including PYY, magainin-2, and skin PYY, a protein homology/analogy recognition engine was used, as reported previously (Kelley and Sternberg, 2009). Primary amino acid sequence alignment was performed with STRAP (Gille et al., 2014).

**Recombinant Peptides**

To study the effect of peptides upon *Candida* *in vitro* and in a murine model of chronic *Candida* challenge *in vivo,* peptides were obtained from GenScript at a purity of >95%. Recombinant peptides were prepared as 2 mM in nuclease-free water (Thermo Fisher Scientific, Ambion, Cat# AM9937), aliquoted, and stored at -80°C. Sequences are as follows:

| **Peptide** | **Amino Acid Sequence** |
| --- | --- |
| Human PYY_1-36_ | YPIKPEAPGEDASPEELNRYYASLRHYLNLVTRQRY |
| Human FITC-PYY_1-36_ | YPI-(FITC-Lys)-PEAPGEDASPEELNRYYASLRHYLNLVTRQRY |
| Human Scrambled PYY_1-36_ | HPYAPLLYTLPQADRSRVLSYEYRNGKEYRAIPENE |
| Magainin-2 | GIGKFLHSAKKFGKAFVGEIMNS |

**Human PYY_1-36_ Bactericidal Assay**

*Lactobacillus rhamnosus* GG*, Enterococcus faecalis, Listeria monocytogenes* EGD*, Peptostreptococcus anaerobius, Staphylococcus aureus* Newman*, Escherichia coli* K12*, Bacteroides fragilis, Salmonella enterica* and *Pseudomonas aeruginosa* were re-inoculated into fresh media and incubated at 37°C to mid-log phase. Cultures were then resuspended in standard assay buffer (10 mM 2-(N-Morpholino)ethanesulfonic acid (MES) pH 6, 25 mM NaCl), pelleted, and diluted 1:25 in standard assay buffer. Recombinant PYY_1-36_ (2 mM) was diluted to concentrations of 10, 20, 30, 40, and 50 *µ*M. Each culture was incubated at 37°C for 2 hours with each PYY concentration. Dilutions of each culture (1:10 and 1:100) and PYY combination were plated on agar plates in duplicate and incubated at 37°C for 24 - 48 hours under aerobic static conditions. CFUs were counted and the average percent of surviving bacteria was calculated based on the CFUs in the 0 µM control for each respective bacterial strain. Three replicates of this experiment were performed and averaged.

**Bacterial Growth Curves**

Overnight cultures of *Enterococcus faecalis* was grown in BHI (230 RPM at 37°C), *Lactobacillus rhamnosus* GG was grown in MRS (static at 37°C), *Listeria monocytogenes* EGD was grown in BHI (230 RPM at 37°C) and *Escherichia coli* K12 was grown in LB (static at 37°C). Cells were passaged into 5 mL of fresh media (1:100) to prepare inocula for growth curves. PYY or PYY derivatives were serially diluted in the appropriate medium in a 96-well plate, for a final concentration of 25, 12.5, 6.25, 3.75, and 1.76 µM. Diluted inoculum (100 µL of) was added to each well, in duplicate, and grown at 37°C statically. OD_600_ was read using a VersaMax plate reader. Bacterial inocula were serially diluted into PBS and plated onto BHI or LB agar plates to determine CFU/well.

***C. albicans* Viability Assay using Propidium Iodide Staining**

To image yeast and hyphal membrane disruption, the DNA binding fluorescent dye, propidium iodide (PI), was utilized. Diluted yeast and hyphae were reconstituted in either RPMI 1640 medium (Thermo Fisher Scientific, Cat# 21870-076), YPD broth (Sigma-Aldrich, Cat# Y1375), RPMI + 2.5% w/v porcine stomach mucin (Sigma-Aldrich, Cat# M11778), standard buffer, or standard buffer + 2.5% w/v porcine stomach mucin. To confirm the localization of PYY, FITC-PYY 1-36 (YPI {Lys (FITC)} PEAPGEDASPEELNRYYASLRHY LNLVTRQRY) at 0, 10, 30, or 60 µM was added to each yeast or hyphae in the various solutions in dark microfuge tubes. Tubes were sealed and incubated for 2 hours at 37°C. PI was then added to each tube and incubated for 5 mins. PYY exposed hyphae or yeast (50 µl) was added to each slide and imaged immediately to detect PI uptake. Leica SP5 STED Laser Scanning Confocal was used for imaging. This experiment was replicated a minimum of 3 times for each variation and a representative image was chosen.

**Growth and Viability Assays for *C. albicans* Yeast**

For yeast form, single colonies were selected and cultured in YPD broth (Sigma-Aldrich, Cat# Y1375) at 30°C overnight with gentle shaking. Cultures were then resuspended in standard buffer/YPD (10 mM MES pH 6, 25 mM NaCl), pelleted (1000 x rcf for 5 minutes at RT) and diluted 1:25 again in standard buffer. Recombinant PYY was added at 37°C for 2 hours. Dilutions at 1:100 were plated on YPD plates and incubated at 30°C for 48 hours. Colonies were counted (CFUs) compared based on the CFUs in the 0 µM control. This was done in duplicate and assayed in triplicate. Yeast growth was measured in a 96-well plate (in triplicate) with varying PYY concentrations, then incubated for 24 hours at 30°C, and O.D. was measured at 600 nm. Due to the varying properties of yeast and hyphae, hyphal viability could not be measured using these methods

***C. albicans* Hyphal Growth Assay**

To induce hyphal growth, overnight yeast cultures were placed into RPMI (Thermo Fisher Scientific, Cat# 21870-076) (1:100) at 37°C for 4 hours. Hyphae cultures (500 µL) were plated (24 well plate) in duplicate with varying PYY concentrations (0, 10, 20, 30, 40, 50, 60, and 100 µM), then incubated at 37°C for 2 hours. The content of each well was resuspended on a pre-weighed 0.8 µm nitrocellulose filter. The filters were left to dry overnight at room temperature, then weighed next day.

**Viability Assay for *C. albicans* Hyphae**

To study the *C. albicans* hyphae viability, the Cell Proliferation Kit II (XTT) (Sigma-Aldrich, Cat# 11465015001) was used. Overnight cultures were diluted 1:100 in RPMI (Millipore Sigma, Cat# R7509, clear medium) or RPMI with 2.5% w/v porcine stomach mucin in a 96-well plate and exposed to various PYY concentrations (0, 10, 20, 30, 40, 50, 60, or 100 µM) in triplicate. Plates were sealed and incubated at 37°C for 2 hours. XTT solution (50 µL) was added to each well according to kit’s protocol, then the plate was incubated at 37°C for an additional hour. The OD was measured with a plate reader at 450 nm and 690 nm as background measurement. These experiments were done in triplicate and replicated 3 times with magainin-2 used as a positive control and PYY scrambled and 0 µM PYY as negative controls.

**Biofilm Assays**

A crystal violet assay was used to assess the effect of PYY on biofilm formation. *C. albicans* hyphae were induced from overnight broth cultures of yeast in RPMI (Thermo Fisher Scientific, Cat# 21870-076) or RPMI + 2.5% w/v mucin from porcine stomach (1:100). Hyphae in RPMI or RPMI + 2.5% w/v mucin were plated into a 96-well plate with varying PYY concentrations (0, 10, 20, 30, 40, 50, 60, or 100 µM) in triplicate and incubated for 24 hours at 37ºC. Biofilms were then gently washed with PBS followed by crystal violet staining, dissolution of the stained biofilm in 30% acetic acid, and quantification of biofilm density by OD_600_ with a plate reader. Magainin-2 was used as a positive control and PYY scrambled and 0 µM as negative controls. Experiments were performed in triplicate with sample triplicates.

**Transmission Electron Microscopy**

Yeast were grown in YPD broth (Sigma-Aldrich, Y1375) overnight at 30ºC. The hyphae forms were then induced by sub-culturing the yeast in RPMI 1640 medium (Thermo Fisher Scientific, 21870-076) at 37ºC for 4 hours without shaking. The yeast and hyphae were harvested by centrifugation at 10,000g x 5 mins, and washed two times with PBS. The cells were then ruptured by bead beating for 2 mins, with 1.0 mm zirconia beads (BioSpec, Bartlesville, OK, 11079). The labeling procedure is adopted from a previous study (Sonnenfeld et al., 1985). Briefly, the ruptured cells were washed two times with 0.05 M (4-(2-Hydroxyethyl)piperazine-1-ethanesulfonic acid (HEPES) buffer (pH 6.8). The ruptured cells were labeled with 0.5 mg/mL cationized ferritin (#15550, Electron Microscopy Services, Hatfield, PA) and dialyzed in 0.05 M HEPES buffer at room for 15 mins. The labeled cells were washed two times with HEPES buffer before they were fixed with 5% glutaraldehyde (in HEPES buffer) for 60 mins. They were then embedded, sectioned and examined by transmission electron microscopy with a FEI Tecnai F30 300kV FEG TEM Electron Microscope (Advanced Electron Microscopy Facility at the University of Chicago).

**Scanning Electron Microscopy**

A single colony of *C. albicans* was inoculated in 5 mL of YPD broth (Sigma-Aldrich, Cat# Y1375) overnight at 30ºC. The hyphal form was then induced by sub-culturing 50 µL of the yeast culture in 5 mL of RPMI 1640 medium (Thermo Fisher Scientific, Cat# 21870-076) at 37ºC for 4 hours without shaking. Yeast and hyphae cultures were inoculated with PYY_1-36_ (30 µM) and H_2_O (control) for 2 hours at 30ºC and 37ºC. Cultures were then spotted on poly-L-lysine-coated coverslips and immersed in 3% glutaraldehyde solution overnight. The coverslips were then washed with ddH_2_O (5 mins x 3) and immersed in 4% OsO_4_ (Sigma-Aldrich, Cat# 11465015001) in ddH_2_O (20 min). The coverslips were dehydrated with the following protocol: 30% ethanol for 5 min, 50% ethanol for 5 min, 70% ethanol for 5 min, 90% ethanol for 5 min, 100% ethanol for 10 min x 2. Fixed coverslips were serial diluted into 100% ethanol and then dried with a Leica EM CPD300 critical point dryer. Dried coverslips were then immediately coated with a 5nm layer of Pt/Pd in a Cressington 208HR sputter coater. Samples were analyzed with a Carl Zeiss Merlin Scanning Electron Microscope (Advanced Electron Microscopy Facility at the University of Chicago).

**Helical Wheel**

The helical wheel was generated with the use of the online tool: heliQuest Analysis (Gautier et al., 2008; *HeliQuest ComputParam form version2*, n.d.). The full PYY_1-36_ sequence was analyzed and the output visualization was used for figure generation. Rotation of the resulting helix was chosen to align vertically with the hydrophobic moment.

**Distal Ileum *Ex vivo* Assay**

Tissues were harvested from PYY-KO and C57Bl6/J WT male mice from 8-10 weeks of age. PYY-KO mice were generated in C57BL/6 blastocysts (Boey et al., 2006). Sections (5 cm) of distal ileum were dissected from the mesentery and ligated on one end. Next, PBS buffer with 1 mM calcium chloride (CaCl_2_) and 1 mM magnesium chloride (MgCl_2_) (pH 7.4), was inoculated with either 2.2 x 10^7^ CFU/mL hyphal form or yeast form of *C. albicans*. Each ileal section was rinsed to remove fecal material and filled and sealed with hyphae, yeast, hyphae supernatant or yeast supernatant and placed in PBS solution at 37°C for 2 hours. Luminal solution, scraped mucus, ileal tissue, and well contents were collected and processed. From each ileal loop, proteins were extracted using cell lysis buffer (1 mL, Cell Signaling Technology, Cat# 9803S) and protease and phosphatase inhibitor (1 µL, Fisher Scientific, Cat# PI78444). Human PYY_1-36_ was spiked in all murine-derived samples as an internal control (10 µM) and samples were processed as described below for mass spectrometry analysis.

**Mass Spectrometry**

Tissue, mucus, lumen and well samples previously prepared by *ex vivo* loop stimulation method were desalted (Pierce Peptide Desalting Columns, Cat# 69570) and lyophilized (n=3). Samples were resuspended in MS-grade H_2_O:MeOH (50:50) and were analyzed by UPLC-ESI-QTOF-MS (Agilent, 6540) with the following gradient (A, 100% H_2_O, 0.1% formic acid; B, 100% MeOH, 0.1% formic acid): 5% B for 2 min, 5-95% B over 8 min, and 95% B for 3 min. Human PYY_1-36_ was spiked in all murine-derived samples as an internal control (10 µM) for normalization and a calibration curve was generated from recombinant human PYY_1-36_ (GenScript). The following *m/z* were observed for murine PYY_1-36_ identification: *m/z* 1414.2 [M+3]^+3^, 707.1 [M+6]^+6^,606.1 [M+7]^+7^. The following *m/z* were observed for human PYY_1-36_: *m/z*, 1437.2 [M+3]^+3^, 1077.9 [M+4]^+4^. The following were searched for murine PYY_3-36_ (M, 3979.00 Da) and PYY_13-36_ (M, 3013.55 Da) with charge states of +1 to +12. Average masses were calculated based on molecular formula with: <https://www.envipat.eawag.ch/index.php>. Analysis was performed with Agilent Mass Hunter (Mass Spectrometry Facility, Department of Chemistry, University of Chicago).

**PYY Partitioning**

A 10 kDa MWCO filter (Thermo Fisher, Cat# PI69570) was placed between porcine mucus (5 mL, pH 7) and ddH_2_O (5 mL, pH 7). Peptides were added on either side and mixed by a rotary shaker for 2 hours at room temperature. The aqueous layer was lyophilized and peptide concentrations measured by mass spectrometry.

**DPP-IV Cleavage of PYY**

Prepared on ice, human PYY_1-36_ (100 µM) was treated with 2.5% w/v porcine mucin (Sigma-Aldrich, Cat# M1778), 50 ng DPP-IV (CedarLane, Cat# CLENZ375) or DPP-IV and mucin for 15 mins at 37ºC in buffer (20 mM Tris, 0.1 M NaCl, 1 mM EDTA, pH 8). DPP-IV was inactivated at 95ºC for 5 mins. Samples were then analyzed by UPLC-ESI-QTOF-MS (Agilent, 6540) and the following ions were quantified: hPYY_1-36_ *m/z* 1077.9 [M+4]^+4^; hPYY_3-36_ *m/z* 1350.5 [M+3]^+3^. Average masses were calculated with: <https://www.envipat.eawag.ch/index.php>

**Caco2 Cell with *C. albicans* and PYY_1-36_ Exposure**

Caco2 cells (ATCC #HTB37) were grown in complete growth medium: 500ml DMED (Fisher cat#10013CV), 200ul transferrin (Gibco BRL 4mg/ml cat#13008-016), 5ml Pen/strep(100x), and 50ml FBS. For passage, Trypsin-EDTA was used to detach the cells; 1:4 or 1:6 sub-cultivation ratio; 1 to 2 times medium renewal per week; routinely tested for mycoplasma contamination. Cells were grown to confluence in 6 well cell culture plates, treated with *C. albicans* (HWP-GFP, which express GFP when hyphal wall protein is expressed) for 6 hours, and fixed with formalin. Cells were rinsed with PBS counter stained with DAPI and imaged in triplicate.

**ITS and 16S rRNA-Based Next Generation Sequencing**

Eleven-week-old PYY KO and WT littermates were used for the microbiome and mycobiome analyses. The ileum and colon were excised from the mice, and luminal contents were collected by a light wash with PBS and mucus from ileum was collected by a gentle scraping. Samples were immediately stored in -80^o^C. Genomic DNA (gDNA) components were extracted using DNeasy PowerSoil Kit (Qiagen, Cat# 12888-100) following the manufacturer's instructions. Samples were sent to Argonne National Laboratory Environmental Sample Preparation and Sequencing Facility (Lemont, IL) for Illumina paired-end Miseq sequencing of the bacterial V4 hypervariable region of the 16S ribosomal RNA (rRNA) gene of bacteria using primers 515F–806R (Caporaso et al., 2011; Parada et al., 2016) and internal transcribed spacer (ITS) of fungi using primers TS1f-ITS2 (Bokulich and Mills, 2013; Kõljalg et al., 2013; White et al., 1990). A total of 74 16S and 74 ITS samples were deposited to the National Centre for Biotechnology Information Sequence Read Archive (accession number SUB7307311). QIIME2 Version 2018.11 (Bolyen et al., 2019) was used to analyze the raw fastq 16S and ITS sequencing data. Divisive Amplicon Denoising Algorithm 2 (DADA2) (Callahan et al., 2016) was used for denoising and quality filtering. Chimeric sequences were removed with default settings. For 16S microbiome analysis, taxonomic classification was performed using Naive Bayes trained q2-feature-classifier 515/506 region from QIIME2 using Greengenes reference database (99% OTUs from 515F/806R region, MD5: 6df67fb01e2f3305e76c61a1c16136b4, 11.2018). For ITS mycobiome analysis, taxonomic classification was performed using UNITE (release date 11.2016) with 99% alignment. Microbiomes with more than 6,000 reads per sample and mycobiomes with more than 400 reads per sample were analyzed and visualized using Calypso (v8.84) (Zakrzewski et al., 2017). Shannon and Simpson diversity indexes were used to determine significance in alpha diversity of the bacterial populations. Chao1 diversity index was used to determine significance in alpha diversity of the fungal community. Principal coordinates analysis (PCoA) using Bray-Curtis and Jaccard distance measures with PERMANOVA was utilized for analyzing bacterial and fungal Beta diversity, respectively. Significance of changes in relative abundance of taxa was assessed by ANOVA. Heatmaps for family and genus levels were generated using Spearman's rank correlation coefficients.

**Intestinal Organoid Studies**

Preparation of Intestinal Epithelial Organoids

Mouse ileal organoids were generated as described (Zhu et al., 2015). Briefly, the small intestine was removed from each mouse, cleaned, and then cut into 1-mm pieces. After multiple washes with ice-cold PBS, intestinal pieces were incubated in 2.5 mM EDTA/PBS, with agitation at 4°C for 60 mins. Cells were then collected in Advanced DMEM/F12 (ADF; Life Technologies, Cat# 12634), and a single-cell suspension passed through a 70-μm cell strainer. Cells were pelleted, then resuspended in 100 μl complete ADF media (ADF supplemented with GlutaMAX; Life Technologies); HEPES buffer (Life Technologies); penicillin and streptomycin (Life Technologies); N2 supplement (Thermo Fisher Scientific, Cat# 17502-048); B-27 Supplement Minus Vitamin A (Thermo Fisher Scientific, Cat# 12587001); murine EGF (50 ng/ml; PeproTech, Cat# 315-09-500ug); 10% Noggin-conditioned media (from 293-Noggin, a generous gift from M. Donowitz, Johns Hopkins University); Jagged-1 (1 μM; Anaspec, Cat# 61298); Y-27632 (10 nM; Cayman Chemical, Cat# 10005583); and R-spondin 1 (500 ng/ml; PeproTech, Cat# 120-38-500ug). Cells were then combined with 200 μl Matrigel (Corning, Cat# cb356239) and plated onto 6-well plates. Matrigel beads containing crypts and cells were allowed to solidify for 1 hour at 5% CO_2_/37°C before adding 2 mL complete ADF media to each well. Complete ADF media were changed every 4 days.

Organoid Imaging

Organoids were collected in an Eppendorf tube and fixed with 4% PFA for 10 mins. They were gently spun down and washed once with 70% ethanol and 2 times with PBS. The primary antibodies (1:200, PYY and lysozyme) were added for 18 hours or overnight in staining buffer (PBS pH 7.4 with 10 mg/mL BSA and 0.3% Triton X-100). The next day, organoids were washed 2 times for 1 hour in PBS and the secondary antibodies were added (1:400) in PBS overnight at 4°C. Again, organoids were washed, 2 times for 1 hour in PBS and counterstained with DAPI for 15 mins (1:1000) in PBS. Following another wash, the organoids were fixed again with 2% PFA in PBS for 10 mins and re-washed. Matrigel mixture (50% Matrigel/50% PBS ~ 40 μl) was added to the tubes containing the organoids, mixed gently, and mounted on the slide. Using a slide pen a thick square was drawn on the slide to contain the organoids. The slides were incubated at 37°C for 1 hour to settle and adhere the organoids and Matrigel mixture to the slide. We used tissue clearing to increase the imaging as described (Lee et al., 2017). Briefly, we incubated the slides for 1 hour each in subsequent mixtures of 20%, 50%, and 80% w/v D-fructose in PBS at 25°C with gentle agitation. We added Prolong Gold and sealed with a coverslip. Imaging was performed under STED (STimulated Emission Depletion) Sp8 confocal microscopy (Leica) at the Integrated Light Microscopy Facility at the University of Chicago.

**QUANTIFICATION AND STATISTICAL ANALYSIS**

GraphPad Prism 8 was utilized to generate all graphs and statistical analysis (<https://www.graphpad.com/scientific-software/prism/>). Figure legends contain statistical analysis and sample size information. Replicates from the *in vitro* assays were normalized to the 0 μM peptide control in each set and combined. Data is presented with mean ± SEM. Statistical significance was analyzed by the Analysis of Variance test (ANOVA) followed by analysis of significance (Dunnett’s Multiple Comparisons Test) or T-test (p<0.05, *; p<0.01, **; p<0.001, ***; p<0.0001, p<0.00001). Data from the *in vivo* studies are presented as mean ± SEM; analyzed with Analysis of Variance test (ANOVA) complemented with (Tukey-Frame’s multiple comparisons test). Data from the *ex vivo* and medium partition studies are presented as mean ± SEM; analyzed with Analysis of Variance test (ANOVA).

For 16S and ITS data, we utilized QIIME2 and Calypso (v8.84). Specification on analysis and sample size can be found in figure legends and ‘Next Generation Sequencing of ITS and 16S’ section in methods.

**KEY RESOURCES TABLE**

| **REAGENT or RESOURCE** | **SOURCE** | **IDENTIFIER** |
| --- | --- | --- |
| Antibodies | | |
| Rabbit polyclonal anti-PYY | abcam | Cat# ab22663, RRID: AB_2175186 |
| Goat polyclonal anti-Lysozyme | Santa Cruz Biotechnology | Cat# C-19 |
| Mouse monoclonal anti-sPLA2 group 2 | Santa Cruz Biotechnology | Cat# SCACC353, RRID: AB_785795 |
| Donkey anti-Rabbit IgG (H+L), Alexa Fluor 555 | Thermo Fisher Scientific | Cat# A-31572, RRID: AB_162543 |
| Donkey anti-Goat IgG (H+L), Alexa Fluor 647 | Thermo Fisher Scientific | Cat# A-21447, RRID: AB_141844 |
| Goat anti-Mouse IgG1, Alexa Fluor 647 | Thermo Fisher Scientific | Cat# A-21240, RRID: AB_141658 |
| Goat polyclonal anti-DPPIV/CD26 | RandD Systems | Cat# AF954, RRID: AB_355739 |
| Bacterial and Virus Strains | | |
| *Lactobacillus rhamnosus GG* | ATCC | Cat# 53103 |
| *Enterococcus faecalis* | ATCC | Cat# BAA-2128 |
| *Listeria monocytogenes* EGD | ATCC | Cat# 49594 |
| *Peptostreptococcus anaerobius* | ATCC | Cat# 27337 |
| *Staphylococcus aureus* Newman | ATCC | Cat# 25904 |
| *Escherichia coli* K12 | ATCC | Cat# PTA-7555 |
| *Bacteroides fragilis* | ATCC | Cat# 25285 |
| *Salmonella enterica* | ATCC | Cat# 1575D-5 |
| *Candida albicans* SC5314 | ATCC | Cat# MYA-2876 |
| Biological Samples | | |
| Human ileal biopsies and tissue sections IRB #15573A | University of Chicago Human Tissue Resource Center | https://htrc.uchicago.edu/ |
| Chemicals, Peptides, and Recombinant Proteins | | |
| Protein Block | Agilent Technologies, DAKO | Cat# X090930-2 |
| Lectin from *Ulex europaeus*, UEA-1, FITC-conjugated | Sigma-Aldrich | Cat# L9006, RRID: AB_2314736 |
| PYY FISH Probe (5’ cy5/ GTGATGGAGTTGGACCAGTG) | IDT DNA | idtdna.com |
| Prolong Gold anti-fade | Thermo Fisher Scientific | Cat# P36930 |
| Recombinant human PYY 1-36 | GenScript | Genscript.com |
| Human FITC-PYY_1-36_ | GenScript | Genscript.com |
| Human scrambled PYY_1-36_ | GenScript | Genscript.com |
| Magainin-2 | GenScript | Genscript.com |
| RPMI-1640 Medium (clear) | MilliporeSigma | Cat# R7509 |
| RPMI-1640 Medium (red) | Thermo Fisher Scientific | Cat# 21870-076 |
| YPD Broth | Sigma-Aldrich | Cat# Y1375 |
| Antibody diluent | Agilent Technologies, DAKO | Cat# S0809 |
| Brain Heart Infusion Broth | Fisher Scientific | Cat# 237500 |
| Brain Heart Infusion Agar | Fisher Scientific | Cat# DF0418-17-7 |
| *Candida* selection plates | BD | Cat# 8012620 |
| MRS Broth | BD Difco | Cat# DF0881175 |
| Porcine stomach mucin | Sigma-Aldrich | Cat# M11778 |
| Luria Broth | BD Difco | Cat# DF0446-07-5 |
| Cationized ferritin | Electron Microscopy Sciences | Cat# 50-980-476 |
| Osmium tetroxide | Sigma-Aldrich | Cat# 75632, CAS# 20816-12-0 |
| Cell lysis buffer (10X) | Cell Signaling Technology | Cat# 9803S |
| Protease phosphatase inhibitor cocktail (100x) | Fisher Scientific | Cat# PI78444 |
| DPP-IV | CedarLane | Cat# CLENZ375 |
| Luria Broth | BD Difco | Cat# DF0446-07-5 |
| Advanced DMEM/F12 | Thermo Fisher Scientific | Cat# 12634 |
| L-Glutamine | Thermo Fisher Scientific | Cat# 25-030-081 |
| N2 Supplement | Thermo Fisher Scientific | Cat# 17502-048 |
| B-27 Supplement Minus Vitamin A | Thermo Fisher Scientific | Cat# 12587001 |
| Murine EGF | PeproTech | Cat# 315-09-500ug |
| Noggin | M. Donowitz, Johns Hopkins University | N/A |
| Jagged-1 | Anaspec | Cat# 61298 |
| Y-27632 | Cayman Chemical | Cat# 10005583, CAS# 129830-38-2 |
| R-Spondin | PeproTech | Cat# 120-38-500ug |
| Matrigel | Corning | Cat# cb356239 |
| Nuclease free water | Thermo Fisher Scientific | Cat# AM9937 |
| Critical Commercial Assays | | |
| Cell Proliferation Kit (XTT) | Sigma-Aldrich | Cat# 11465015001 |
| DNeasy PowerSoil Kit | Qiagen | Cat# 12888-100 |
| Rneasy Micro Kit | Qiagen | Cat# 74004 |
| Deposited Data | | |
| ITS and 16SrRNA Data | This paper | National Centre for Biotechnology Information Sequence Read Archive: # SUB7307311 |
| Single cell RNAseq Analysis | Previously published | Haber, et al., 2017 |
| Experimental Models: Cell Lines | | |
| Caco2 (HTB-37) | ATCC | <https://www.atcc.org/products/all/htb-37.aspx> |
| Human-derived organoids (IRB Approval# IRB #15573A, UChicago) | Tissue Engineering and Cell Models Core, DDRCC, UChicago | |
| Experimental Models: Organisms/Strains | | |
| *Candida albicans* HGFP3 | UChicago, Dr. John Alverdy, (Chairatana and Nolan, 2017) | N/A |
| *Pseudomonas aeruginosa* MPAO1-P2 | UChicago, Dr. John Alverdy,(Luong et al., 2014) | N/A |
| C57BL/6J | The Jackson Laboratory | <https://www.jax.org/strain/000664> |
| PYY^-/-^ gene-deficient mice on the C57Bl/6 background | Herber Herzog (Boey et al., 2006) | N/A |
| Oligonucleotides | | |
| Primer: PYY, Forward -GTTAACTACACCGACTTCACT: Reverse- GTCCGAGACACCGAGATA | This study | idtdna.com |
| Primer: Lysozyme (Lyz1), Forward - ATAAATTCTCAGCTCATGTGTC: Reverse- TCTTCTGTCTCAGAGTTAGTATG | This study | idtdna.com |
| Primer: Cryptdin 1, Forward - AAGAGACTAAAACTGAGGAGC: Reverse- GGACACCAGTACTCACATTC | This study | idtdna.com |
| Primer: Sucrase Isomaltase, Forward - TATGGTGGGAATGAACAACAG: Reverse- CAGGAATTCAAACTGATGGTG | This study | idtdna.com |
| Recombinant DNA | | |
| N/A |  |  |
| Software and Algorithms | | |
| enviPat Web 2.4 | Loos et al., 2015 | <https://www.envipat.eawag.ch/index.php> |
| Agilent Mass Hunter | Agilent Technologies | <https://www.agilent.com/en/products/software-informatics/mass-spectrometry-software> |
| GraphPad Prism 8 | Prism 8, graphpad.com | <https://www.graphpad.com/scientific-software/prism/> |
| Image Pro-Plus Software | Media Cybernetics | <https://www.mediacy.com/imageproplus> |
| ThunderSTORM, ImageJ Plug-In | Ovesný et al., 2014 | <https://doi.org/10.1093/bioinformatics/btu202> |
| Pep-Calc | Lear and Cobb, 2016 | <https://www.pep-calc.com/> |
| STRAP | Gille et al., 2014 | <https://doi.org/10.1093/nar/gku400> |
| ImageJ 1.x | Schneider et al., 2012 | https://doi.org/10.1038/nmeth.2089 |
| heliQuest | Gautier et al., 2008 | <https://heliquest.ipmc.cnrs.fr/> |
| QIIME2 2018.11 | Bolyen et al., 2019 | https://doi.org/10.1038/s41587-019-0209-9 |
| Calypso (v8.84) | Zakrzewski et al., 2017 | <https://doi.org/10.1093/bioinformatics/btw725> |
| UNITE | Release date 11.2016.(Kõljalg et al., 2013) | <https://unite.ut.ee/repository.php> |
| DADA2 | Callahan et al., 2016 | QIIME2 Plugin |
| Greengenes | Release data 11.2018. | <http://greengenes.secondgenome.com/> |

**Materials and Methods References**

Batterham, R. L., Heffron, H., Kapoor, S., Chivers, J. E., Chandarana, K., Herzog, H., Le Roux, C. W., Thomas, E. L., Bell, J. D., & Withers, D. J. (2006). Critical role for peptide YY in protein-mediated satiation and body-weight regulation. *Cell Metabolism*, *4*(3), 223–233. https://doi.org/10.1016/j.cmet.2006.08.001

Boey, D., Lin, S., Karl, T., Baldock, P., Lee, N., Enriquez, R., Couzens, M., Slack, K., Dallmann, R., Sainsbury, A., and Herzog, H. (2006). Peptide YY ablation in mice leads to the development of hyperinsulinemia and obesity. *Diabetologia*, *49*(6), 1360–1370. https://doi.org/10.1007/s00125-006-0237-0

Bokulich, N. A., and Mills, D. A. (2013). Improved selection of internal transcribed spacer-specific primers enables quantitative, ultra-high-throughput profiling of fungal communities. *Applied and Environmental Microbiology*, *79*(8), 2519–2526. https://doi.org/10.1128/AEM.03870-12

Bolyen, E., Rideout, J. R., Dillon, M. R., Bokulich, N. A., Abnet, C. C., Al-Ghalith, G. A., Alexander, H., Alm, E. J., Arumugam, M., Asnicar, F., Bai, Y., Bisanz, J. E., Bittinger, K., Brejnrod, A., Brislawn, C. J., Brown, C. T., Callahan, B. J., Caraballo-Rodríguez, A. M., Chase, J., … Caporaso, J. G. (2019). Reproducible, interactive, scalable and extensible microbiome data science using QIIME 2. In *Nature Biotechnology* (Vol. 37, Issue 8, pp. 852–857). Nature Publishing Group. https://doi.org/10.1038/s41587-019-0209-9

Callahan, B. J., McMurdie, P. J., Rosen, M. J., Han, A. W., Johnson, A. J. A., and Holmes, S. P. (2016). DADA2: High-resolution sample inference from Illumina amplicon data. *Nature Methods*, *13*(7), 581–583. https://doi.org/10.1038/nmeth.3869

Caporaso, J. G., Lauber, C. L., Walters, W. A., Berg-Lyons, D., Lozupone, C. A., Turnbaugh, P. J., Fierer, N., and Knight, R. (2011). Global patterns of 16S rRNA diversity at a depth of millions of sequences per sample. *Proceedings of the National Academy of Sciences of the United States of America*, *108*(SUPPL. 1), 4516–4522. https://doi.org/10.1073/pnas.1000080107

Chairatana, P., and Nolan, E. M. (2017). Human α-Defensin 6: A Small Peptide That Self-Assembles and Protects the Host by Entangling Microbes. *Accounts of Chemical Research*, *50*(4), 960–967. https://doi.org/10.1021/acs.accounts.6b00653

Gautier, R., Douguet, D., Antonny, B., and Drin, G. (2008). HELIQUEST: A web server to screen sequences with specific α-helical properties. *Bioinformatics*, *24*(18), 2101–2102. https://doi.org/10.1093/bioinformatics/btn392

Gille, C., Fähling, M., Weyand, B., Wieland, T., and Gille, A. (2014). Alignment-Annotator web server: Rendering and annotating sequence alignments. *Nucleic Acids Research*, *42*(W1). https://doi.org/10.1093/nar/gku400

Haber, A. L., Biton, M., Rogel, N., Herbst, R. H., Shekhar, K., Smillie, C., Burgin, G., Delorey, T. M., Howitt, M. R., Katz, Y., Tirosh, I., Beyaz, S., Dionne, D., Zhang, M., Raychowdhury, R., Garrett, W. S., Rozenblatt-Rosen, O., Shi, H. N., Yilmaz, O., … Regev, A. (2017). A single-cell survey of the small intestinal epithelium. *Nature*, *551*(7680), 333–339. https://doi.org/10.1038/nature24489

*HeliQuest ComputParam form version2*. (n.d.). Retrieved April 27, 2020, from https://heliquest.ipmc.cnrs.fr/cgi-bin/ComputParams.py

Kelley, L. A., and Sternberg, M. J. E. (2009). Protein structure prediction on the web: A case study using the phyre server. *Nature Protocols*, *4*(3), 363–373. https://doi.org/10.1038/nprot.2009.2

Kõljalg, U., Nilsson, R. H., Abarenkov, K., Tedersoo, L., Taylor, A. F. S., Bahram, M., Bates, S. T., Bruns, T. D., Bengtsson-Palme, J., Callaghan, T. M., Douglas, B., Drenkhan, T., Eberhardt, U., Dueñas, M., Grebenc, T., Griffith, G. W., Hartmann, M., Kirk, P. M., Kohout, P., … Larsson, K. H. (2013). Towards a unified paradigm for sequence-based identification of fungi. In *Molecular Ecology* (Vol. 22, Issue 21, pp. 5271–5277). John Wiley and Sons, Ltd. https://doi.org/10.1111/mec.12481

Lear, S., and Cobb, S. L. (2016). Pep-Calc.com: A set of web utilities for the calculation of peptide and peptoid properties and automatic mass spectral peak assignment. *Journal of Computer-Aided Molecular Design*, *30*(3), 271–277. https://doi.org/10.1007/s10822-016-9902-7

Lee, S. S. Y., Bindokas, V. P., and Kron, S. J. (2017). Multiplex three-dimensional optical mapping of tumor immune microenvironment. *Scientific Reports*, *7*(1), 1–11. https://doi.org/10.1038/s41598-017-16987-x

Loos, M., Gerber, C., Corona, F., Hollender, J., and Singer, H. (2015). Accelerated isotope fine structure calculation using pruned transition trees. *Analytical Chemistry*, *87*(11), 5738–5744. https://doi.org/10.1021/acs.analchem.5b00941

Luong, P. M., Shogan, B. D., Zaborin, A., Belogortseva, N., Shrout, J. D., Zaborina, O., and Alverdy, J. C. (2014). Emergence of the P2 Phenotype in Pseudomonas aeruginosa PAO1 Strains Involves Various Mutations in mexT or mexF. *Journal of Bacteriology*, *196*(2), 504–513. https://doi.org/10.1128/JB.01050-13

Ovesný, M., Křížek, P., Borkovec, J., Švindrych, Z., and Hagen, G. M. (2014). ThunderSTORM: A comprehensive ImageJ plug-in for PALM and STORM data analysis and super-resolution imaging. *Bioinformatics*, *30*(16), 2389–2390. https://doi.org/10.1093/bioinformatics/btu202

Parada, A. E., Needham, D. M., and Fuhrman, J. A. (2016). Every base matters: assessing small subunit rRNA primers for marine microbiomes with mock communities, time series and global field samples. *Environmental Microbiology*, *18*(5), 1403–1414. https://doi.org/10.1111/1462-2920.13023

Schneider, C. A., Rasband, W. S., and Eliceiri, K. W. (2012). NIH Image to ImageJ: 25 years of image analysis. *Nature Methods*, *9*(7), 671–675. https://doi.org/10.1038/nmeth.2089

Sonnenfeld, E. M., Beveridge, T. J., and Doyle, R. J. (1985). Discontinuity of charge on cell wall poles of Bacillus subtilis. *Canadian Journal of Microbiology*, *31*(9), 875–877. https://doi.org/10.1139/m85-163

White, T. J., Bruns, T., Lee, S., and Taylor, J. (1990). Amplification and direct sequencing of fungal ribosomal RNA genes for phylogenetics. In *PCR Protocols: A Guide to Methods and Applications* (pp. 315–322). Elsevier. https://doi.org/10.1016/B978-0-12-372180-8.50042-1

Zakrzewski, M., Proietti, C., Ellis, J. J., Hasan, S., Brion, M. J., Berger, B., and Krause, L. (2017). Calypso: A user-friendly web-server for mining and visualizing microbiome-environment interactions. *Bioinformatics*, *33*(5), 782–783. https://doi.org/10.1093/bioinformatics/btw725

Zhu, X., Messer, J. S., Wang, Y., Lin, F., Cham, C. M., Chang, J., Billiar, T. R., Lotze, M. T., Boone, D. L., and Chang, E. B. (2015). Cytosolic HMGB1 controls the cellular autophagy/apoptosis checkpoint during inflammation. *Journal of Clinical Investigation*, *125*(3), 1098–1110. https://doi.org/10.1172/JCI76344
